# Aberrant association of chromatin with nuclear periphery induced by Rif1 leads to mitotic defect and cell death

**DOI:** 10.1101/2022.05.13.491604

**Authors:** Yutaka Kanoh, Seiji Matsumoto, Masaru Ueno, Motoshi Hayano, Satomi Kudo, Hisao Masai

## Abstract

Chromatin is compartmentalized in nuclei and its architecture and nuclear location may have impacts on chromatin events. Rif1, identified as a potent suppressor of *hsk1-null* mutation (defective in initiation of DNA replication) in fission yeast, recognizes G-quadruplex structures and inhibits origin firing in their 50∼100-kb vicinity, leading us to postulate that Rif1 may generate chromatin higher-order structures inhibitory for initiation. However, effects of Rif1 on chromatin localization in nuclei have not been known. We show here that overexpression of Rif1 causes growth inhibition and eventually cell death in fission yeast. Chromatin binding activities of Rif1, but not recruitment of phosphatase PP1, are required for growth inhibitory effect. Overexpression of a PP1 binding site mutant of Rif1 does not delay S-phase, but still causes cell death, indicating that cell death is caused not by S-phase problems but by issues in other phases of cell cycle, most likely M-phase. Indeed, Rif1 overexpression generates cells with unequally segregated chromosomes. Rif1 overexpression relocates chromatin near nuclear periphery in a manner dependent on its chromatin-binding ability, and this correlates with growth inhibition and cell death induction. Thus, regulated Rif1-mediated chromatin association with nuclear periphery is important for coordinated progression of S- and M-phases.

## Introduction

Chromosomes are packaged and compartmentalized in nuclei in an organized manner and their spatial arrangement and regional interactions regulate various chromosome transactions. Presence of chromosome near nuclear periphery is generally associated with inactive transcription or late replication (Lemaitre & Bickmore, 2015; Mahamid et al., 2016). The nuclear lamina, nuclear pore complexes and epigenetic regulators have been implicated in regulating tethering of chromatin at nuclear periphery (Amiad-Pavlov et al., 2021; Arib & Akhtar, 2011; Laghmach, Di Pierro, & Potoyan, 2021; Ptak & Wozniak, 2016; See et al., 2020; Smith, Poleshko, & Epstein, 2021). Rif1 was identified as a regulator of replication timing and was shown to suppress late origin firing. On the basis of effect of a Rif1 binding site mutant on origin firing patterns and biochemical functions of Rif1 protein, we have proposed that Rif1 regulates replication timing partly through generating specific chromatin architecture (Y. Kanoh et al., 2015) (Kobayashi et al., 2019; Masai et al., 2019). In mammalian cells, Rif1 is localized near nuclear periphery and was shown to regulate chromatin loop formation (Yamazaki et al., 2012). However, how Rif1 regulates chromatin localization and its potential roles in regulation of cellular events have not been known. We have tested this by examining the effects of overexpression of Rif1 on the cell cycle progression and chromatin states in fission yeast cells (Cornacchia et al., 2012; Foti et al., 2016; Yamazaki et al., 2012).

Rif1, originally isolated as a telomere binding factor, is a multifunctional protein that regulates various aspects of chromosome dynamics, including DSB repair, DNA replication, recombination, transcription and others (Campolo et al., 2013; Cornacchia et al., 2012; Di Virgilio et al., 2013; Hayano et al., 2012; Klein et al., 2021; Yamazaki et al., 2012; Zimmermann, Lottersberger, Buonomo, Sfeir, & de Lange, 2013). Fission yeast Rif1 binds to chromatin, most notably at the telomere and telomeres are elongated in *rif1*Δ cells (J. Kanoh & Ishikawa, 2001; Kobayashi et al., 2019). Hayano *et al*. reported that *rif1*Δ can restore the growth of *hsk1*Δ cells (Hayano et al., 2012), indicating that the loss of *rif1* can bypass the requirement for the fission yeast “Cdc7” kinase (Dbf4-dependent kinase; DDK), which is essential for replication initiation under normal growth conditions. Late-firing origins are extensively deregulated in *rif1*Δ cells, consistent with a role for Rif1 in suppressing late-firing origins. Similarly, mammalian Rif1 was also found to regulate the genome-wide replication timing (Cornacchia et al., 2012; Klein et al., 2021; Yamazaki et al., 2012). Rif1 interacts with PP1 phosphatase through its PP1 binding motifs, and the recruitment of the phosphatase by Rif1 counteracts the phosphorylation events that are essential for initiation of DNA replication including the phosphorylation of Mcm, explaining a mechanism of Rif1-mediated inhibition of replication initiation (Dave, Cooley, Garg, & Bianchi, 2014; Hiraga et al., 2014; Mattarocci et al., 2014; Shyian et al., 2016).

In addition to its binding to telomeres, fission yeast Rif1 also binds to the arm segments of the chromosomes. Thirty-five strong Rif1 binding sites (Rif1bs) have been identified on fission yeast chromosomes. These sequences contain multiple G-tracts and have propensity to form G-quadruplex (G4) structures (Y. Kanoh et al., 2015).

Consistent with this, Rif1 specifically binds to G4-containing DNA *in vitro*, and mutations of G-tracts impaired both *in vivo* chromatin binding of Rif1 and *in vitro* interaction of Rif1 with the Rif1bs. Notably, loss of Rif1 binding at a single Rif1bs caused deregulation of late firing origins in the 50∼100kb segment in its vicinity, consistent with the notion that Rif1 binding generates a chromosome compartment where origin firings are suppressed.

In mammals, Rif1 is preferentially localized at the nuclear periphery in the Triton X-100- and DNase I-resistant compartments, where it regulates the length of chromatin loops (Yamazaki et al., 2012). In fission yeast, Rif1 is biochemically fractionated into Triton X-100- and DNase I-insoluble fractions(Y. Kanoh et al., 2015).

In budding yeast, Rif1 was shown to be palmitoylated and the lipid modification-mediated membrane association plays important roles in DSB repair (Fontana et al., 2019; Park et al., 2011). However, it is unknown whether similar mechanisms operate for Rif1 from other species.

We hypothesized that Rif1 generates higher-order chromatin architecture through its ability to tether chromatin loops, and that this chromatin structure constitutes the replication inhibitory chromatin compartments that are deregulated during mid-S phase. In order to gain more insight into the roles of Rif1 in regulation of chromatin structure and cell cycle, we have analyzed the effect of Rif1 overexpression on the growth, cell cycle progression and chromatin structure in fission yeast. The results indicate that association of chromatin with nuclear periphery needs to be precisely regulated for proper S and M phase progression.

## Results

### Overexpression of Rif1 prevents cell growth

The *rif1* mutation was identified as a suppressor of the *hsk1-null* mutation (encoding Cdc7 kinase [DDK] homologue) in fission yeast (Hayano et al., 2012) and we previously showed that Rif1 suppressed firing of origins over ∼100kb segments spanning its binding sites (Y. Kanoh et al., 2015). During the course of our experiments, we cloned the *rif1^+^* ORF into pREP41 and expressed Rif1 under the inducible nmt41 (<u>N</u>o <u>M</u>essage in <u>T</u>hiamine 1) promoter (**Fig. 1A and B**). Induction of the full length Rif1 (1-1400aa) in medium without thiamine strongly inhibited growth of both *hsk1^+^* and *hsk1-89* (a temperature-sensitive stain) cells (**Fig. 1C and D**). Various truncated deletion mutants of Rif1 were cloned and expressed from the inducible nmt41 promoter to examine growth in *hsk1^+^*or *hsk1-89* cells (**Fig. 1B**). The expression levels of these truncation mutants were examined by western blotting and the results indicate that all the mutants are uniformly expressed (Supplementary data **Fig. S1**). A C-terminal 140-amino acids deletion (construct 1-1260aa) inhibited the growth of *hsk1^+^* weakly and *hsk1-89* strongly (**Fig. 1C and D**). Further truncations of the Rif1 C-terminus (constructs 1-965aa and 1-442aa) result in complete loss of growth inhibition. We previously showed that truncation of the Rif1 C-terminal 140-amino acids (construct 1-1260aa) resulted in loss of telomere length regulation (Kobayashi et al., 2019). Therefore, we conclude that the growth inhibition caused by overexpression of Rif1 does not depend on its function in telomere regulation. In contrast, deletion of the N-terminal 150-amino acids from Rif1 resulted in loss of growth inhibition (**Fig. 1C and D**). However, deletion of the N-terminal 80-amino acids did not affect the ability of Rif1 to inhibit the growth in the *hsk1^+^* cells (**Fig. 1C**). The N-terminal domain (88-1023aa) of fission yeast Rif1 is predicted to form a 3-D structure identical to that of the HEAT repeats by AlphaFold2 (Jumper et al., 2021). Thus, it is highly likely that the HEAT domain is required for growth inhibition by Rif1.

**Figure 1.**
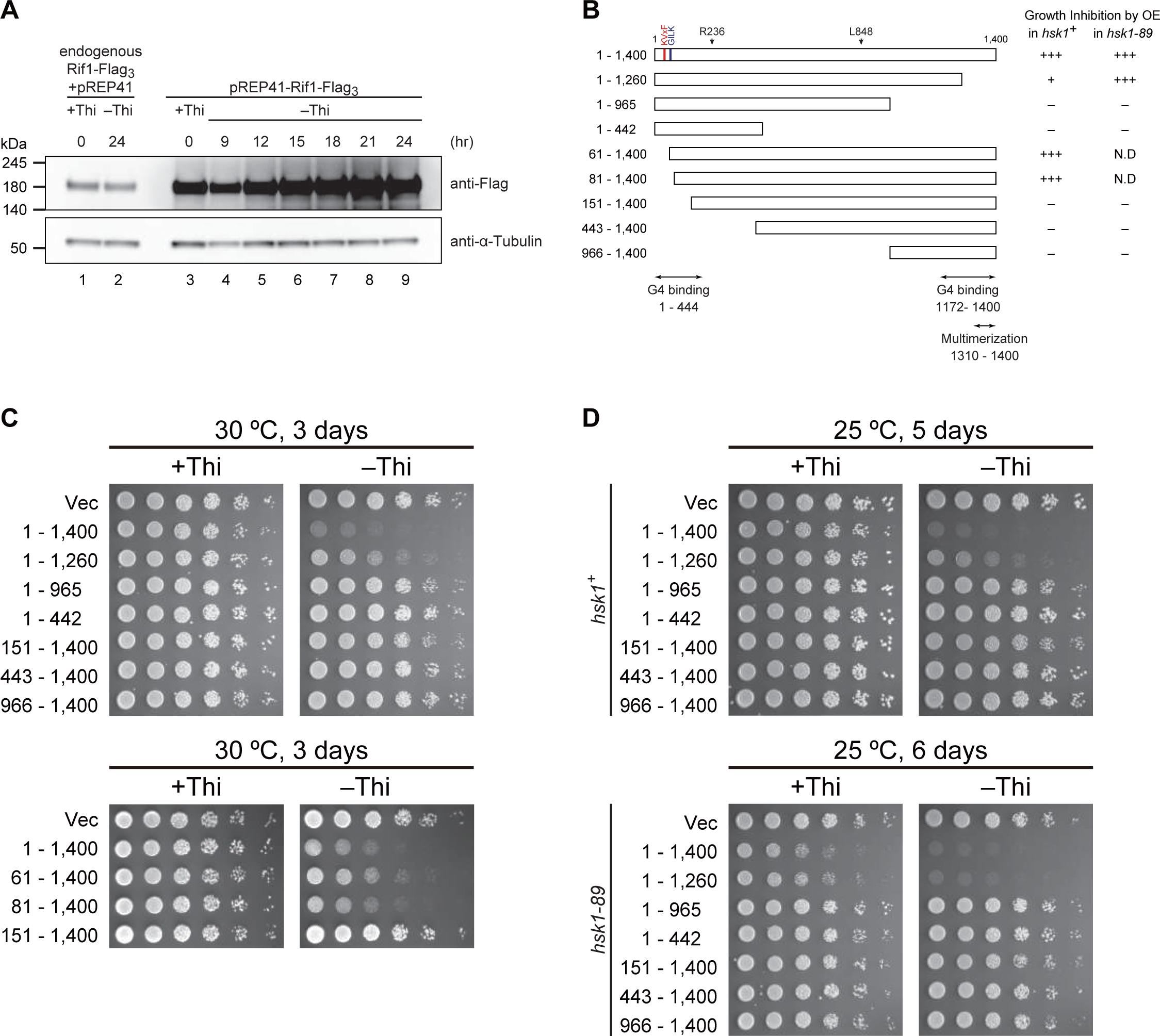
Overexpression of Rif1 inhibits growth of fission yeast cells. **(A)** Time course of overexpression of Rif1-Flag_3_ protein expressed on pREP41 plasmid under the nmt41promoter after transfer to medium lacking thiamine (lanes 3-9). Lanes 1 and 2, Rif1-Flag_3_ is expressed at the endogenous *rif1* locus under its own promoter in the presence or absence of thiamine in the medium. Proteins were detected by the anti-Flag antibody. (**B)** Schematic drawing of deletion derivatives of Rif1 protein analyzed in this study. + indicates growth inhibition, whereas – indicates the absence of growth inhibition. The PP1 binding motifs (RVxF and SILK) are indicated in red and blue, respectively. Note that the motifs in Rif1 are slightly diverged from the above consensus sequences. The polypeptide segments capable of G4 binding and oligomerization are also indicated. (**C**-**D**) Effects of overexpression of the full length and truncated mutants of Rif1 were examined. Proteins were expressed on pREP41 in medium containing (+Thi) or lacking (–Thi) thiamine. Serially diluted (5× fold) cells were spotted and growth of the spotted cells was examined after incubation at the indicated temperature for the indicated time. Growth inhibition was observed with full-length (1-1400), 1-1260, 61-1400 or 81-1400 derivatives of Rif1.

### Growth inhibition by Rif1-overexpression does not involve recruitment of PP1

Rif1 recruits protein phosphatase 1 (PP1) through its PP1 binding motifs (Rif1_40-43_ and Rif1_64-67_) and the recruited PP1 counteracts the phosphorylation by Cdc7 kinase at origins (Dave et al., 2014; Hiraga et al., 2014; Mattarocci et al., 2014). This interaction of Rif1 with PP1 is crucial for replication inhibition by Rif1 at late origins. Therefore, we examined whether Rif1-overexpression-induced growth inhibition is caused by hyper-recruitment of PP1. Fission yeast cells have two PP1 genes *dis2^+^* and *sds21^+^*. A single disruption mutation of *dis2* or *sds21* is viable, but the double mutation is lethal (Kinoshita, Ohkura, & Yanagida, 1990). *dis2-11* is a cold-sensitive mutant of *dis2^+^*. We first examined whether growth inhibition caused by Rif1 overexpression depends on the PP1 genes. Overexpression of Rif1 in *dis2-11*, *dis2*Δ and *sds21*Δ resulted in strong growth inhibition in all the tested strains on EMM media without thiamine (**Fig. 2A**). The extent of inhibition in each PP1 mutant was as strong as that observed in the wild-type, suggesting that the recruited PP1 is not responsible for the growth inhibition.

**Figure 2.**
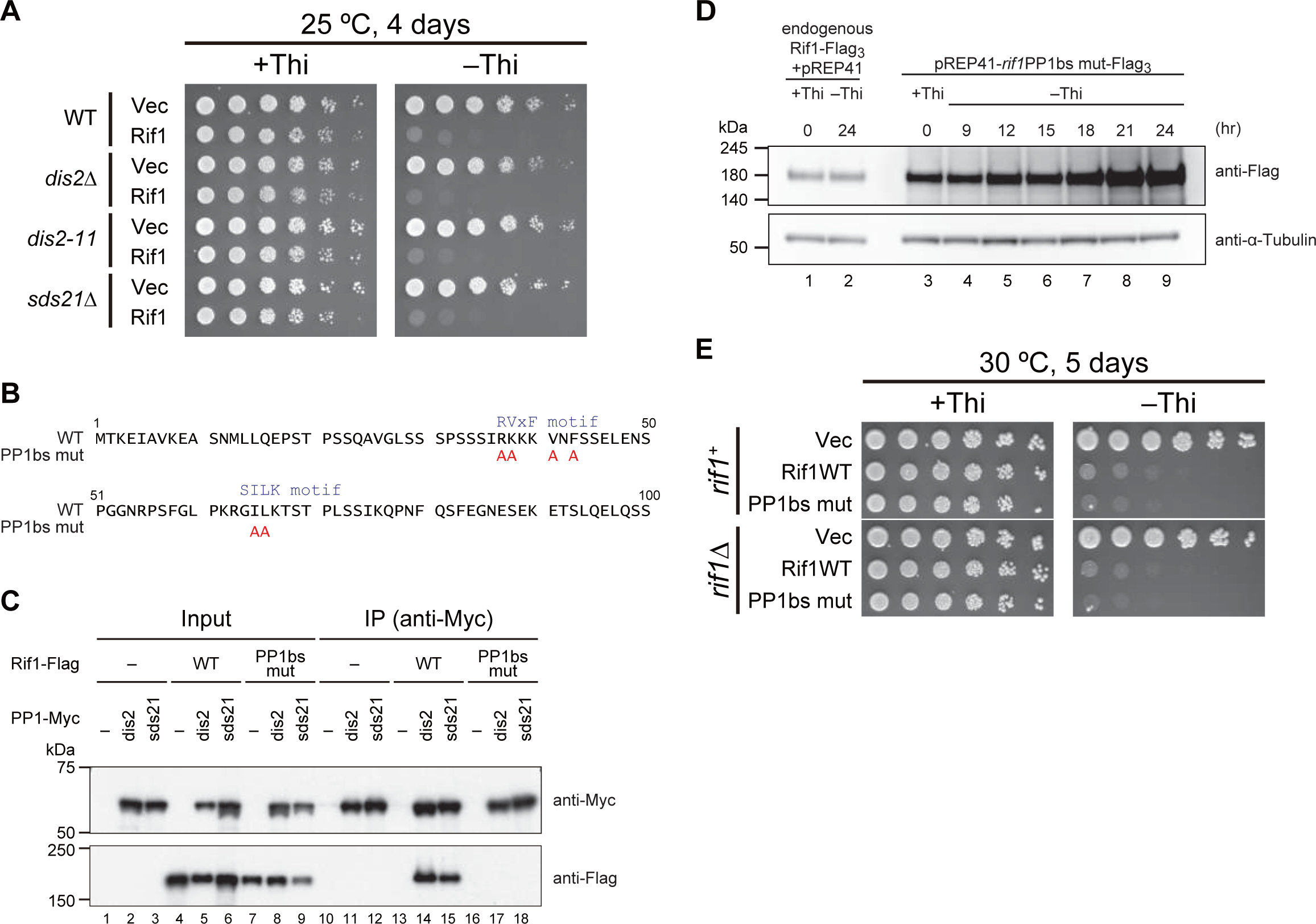
PP1-Rif1 interaction is not required for the growth inhibition caused by Rif1 overexpression. (**A**) Spot tests of Rif1 overexpression in the PP1 mutants *dis2*Δ, *dis2-11* or *sds21*Δ cells were conducted as described in Fig. 1C and **D**. Rif1 overexpression inhibited growth of the mutant cells similar to the wild type cells. (**B**) Mutations introduced at the PP1 binding sites (RVxF and SILK motif) of Rif1. (**C**) Using the extracts made from the cells expressing both Flag-tagged Rif1 and Myc-tagged PP1 (Dis2 or Sds21), PP1 were immunoprecipitated by anti-Myc antibody, and co-immunoprecipitated Rif1 were detected. The PP1bs mutant of Rif1 does not interact with either PP1. (**D**) Time course of overexpression of *rif1*PP1bsmut-Flag_3_ protein expressed on pREP41 plasmid under the nmt41 promoter after transfer to medium lacking thiamine (lanes 3-9). Lanes 1 and 2, *rif1*PP1bsmut -Flag_3_ is expressed at the endogenous *rif1* locus under its own promoter in the presence or absence of thiamine in the medium. (**E**) Spot tests of the wild-type and *rif1*Δ cells overexpressing the wild-type or a PP1bs mutant. Overexpression of the PP1bs mutant Rif1 inhibited growth of fission yeast cells in a manner similar to or slightly better than the wild-type Rif1 did.

To further examine the involvement of PP1, we constructed a PP1-binding mutant of Rif1. We generated an alanine-substituted mutant of the two PP1 binding motifs (KVxF at aa40-43 and GILK at aa64-67) of Rif1 (**Fig. 2B**) and confirmed, by immunoprecipitation, that the mutant Rif1 (PP1bsmut) did not bind to either Dis2 or Sds21 (**Fig. 2C**; compare lanes14 and 17, lanes 15 and 18). Growth inhibition caused by overexpression of the PP1bsmut was comparable to or even slightly stronger than that caused by the wild-type Rif1 in both *rif1*^+^ and *rif1*Δ backgrounds (**Fig. 2D and E**). This is a further support for the conclusion that growth inhibition by Rif1-overexpression does not depend on recruitment of PP1. These results are consistent with the observation that the N-terminal truncation Rif1 (81-1400aa), which lacks the PP1 binding sites, can inhibit the growth upon overexpression (**Fig. 1C**).

### Growth inhibition by Rif1-overexpression does not depend on Taz1 or replication checkpoint

We next asked whether the growth inhibition by overexpressed Rif1 is caused by counteracting the Hsk1-Dfp1/Him1 activity. Coexpression of both Hsk1 and Dfp1/Him1 under the control of the nmt1 promoter itself caused growth inhibition in fission yeast cells, and growth was partially restored by *rif1* deletion. However, overexpression of Hsk1-Dfp1/Him1 did not improve the growth of Rif1-overexpressing cells and inhibited the growth more strongly (**Supplementary data Fig. S2A**), excluding the possibility that inhibition of growth is due to the reduced Hsk1 kinase actions. Consistent with the results of the C-terminal deletion mutant (1-1260aa), which does not interact with Taz1 but still inhibits growth (**Fig. 1C and D**), mutation of *taz1^+^* known to be required for telomere-localization of Rif1 did not affect the growth inhibition by Rif1-overexpression (**Supplementary data Fig. S2B**). Similarly, growth inhibition was observed in mutants of the replication checkpoint genes (Furuya & Carr, 2003), *rad3 tel1*, *rad3*, *chk1* or *cds1* (**Supplementary data Fig. S2B**). The extent of growth inhibition was not affected by *cdc25-22* or *wee1-50*, genes involved in mitotic checkpoint (Iino & Yamamoto, 1997; Kumar & Huberman, 2004; Rowley, Hudson, & Young, 1992) (**Supplementary data Fig. S2C**). These results suggest that the growth inhibition is not caused by replication or mitotic checkpoint functions.

### Effect of Rif1 overexpression on entry into S phase and replication checkpoint activation

In order to clarify the mechanisms of Rif1-mediated growth inhibition, we examined whether Rif1 overexpression inhibits S phase initiation and progression. We synchronized cell cycle by release from *nda3*-mediated M phase arrest, and analyzed the DNA content by FACS. In the wild-type cells, DNA synthesis was observed at 18.5 hr from the release, and continued until 19.5 hr. In Pnmt1-Rif1, where Rif1 was overexpressed, DNA synthesis was delayed by 30 min and was not completed even at 20 hr, indicating that Rif1 overexpression retarded the initiation and elongation of DNA synthesis. In contrast, in Pnmt1-*rif1*PP1bsmut, DNA synthesis occurred with timing similar to the wild-type, indicating that the overexpression of the PP1 mutant Rif1 does not affect the S phase (**Fig. 3A and B**). This suggests that inhibition of S phase by overexpressed Rif1 is due to the hyper-recruitment of PP1, which would counteract the phosphorylation events mediated by Cdc7 or Cdk and inhibit initiation.

**Figure 3.**
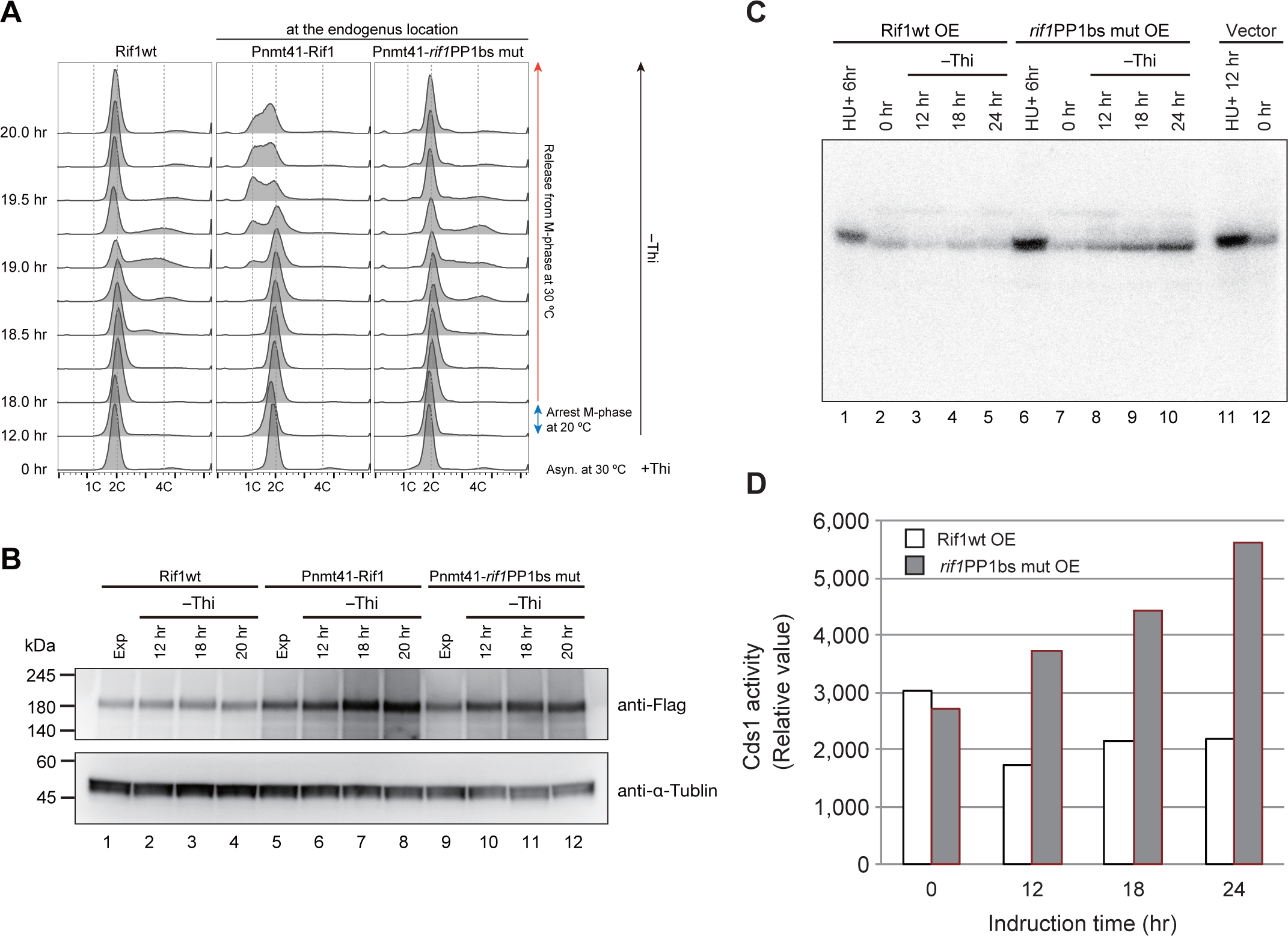
Effect of overexpression of Rif1 protein on cell cycle progression and replication checkpoint activation. (**A**) The *nda3-KM311* cold-sensitive mutant cells with wild-type *rif1*^+^ or those expressing the wild-type Rif1 (Pnmt1-Rif1) or PP1bs mutant Rif1 (Pnmt1-*rif1*PP1bsmut) at the endogenous *rif1* locus under nmt1 promoter were arrested at M-phase by incubation at 20°C for 6 hr with concomitant depletion of thiamine. The cells were released into cell cycle at 30°C. The cell cycle progression was monitored by flow cytometry. The cells with Pnmt1-*rif1*PP1bsmut entered S-phase at 30 min after release, similar to the *rif1*^+^ cells, whereas those with Pnmt1-Rif1 entered S-phase later (>60 min after release). (**B**) The level of Rif1 in the samples from **A** was examined by western blotting. (**C**) The cells harboring Rif1 (wt or *rif1*PP1bsmut)-expressing plasmid or vector, as indicated, were starved for thiamine for the time indicated. The whole cell extracts were prepared and were run on SDS-PAGE containing MBP in the gel. In-gel kinase assays were conducted as described in “Materials and Methods”. HU, treated with 2 mM HU for the time indicated as a positive control of Cds1 activation. (**D**) Quantification of the results in **C**.

We next examined whether overexpression of Rif1 activates replication checkpoint. We measured Cds1 kinase activity by in-gel kinase assay. While Cds1 kinase activity decreased at 6 hr after induction and then slightly increased afterward in cells overexpressing the wild-type Rif1, it continued to increase until 24 hr after overexpression of the PP1bs mutant. These results indicate that overexpression of the Rif1 PP1bs mutant activates replication checkpoint, whereas the wild-type Rif1 does not, consistent with the above results that growth inhibition is not caused by replication checkpoint.

### Short spindles and abnormally segregated nuclei are accumulated in cells **overexpressing Rif1.**

We next observed the morphological effects of Rif1-overexpression in cells expressing GFP-tagged histone H3 (*hht2+-GFP*) or GFP-tagged αtubulin (*GFP-α2tub*). Cut cells appeared in Rif1-overexpressing cells, indicative of failure of chromosome segregation (**Fig. 4A**). At 22 hr after induction, cells with abnormal nuclei accumulated and, notably, cells with unequally segregated nuclei reached more than 23% of the cell population (**Fig. 4B**). We then examined whether DNA damages are induced in these cells by measuring the cells with Rad52 foci, an indicator of DSB (Du, Nakamura, Moser, & Russell, 2003; Matsumoto, Ogino, Noguchi, Russell, & Masai, 2005). Cells with nuclei containing Rad52 foci accumulated in Rif1-overexpressing cells (up to 44 % of the cells at 72 hr after induction; **Fig. 4C**).

**Figure 4.**
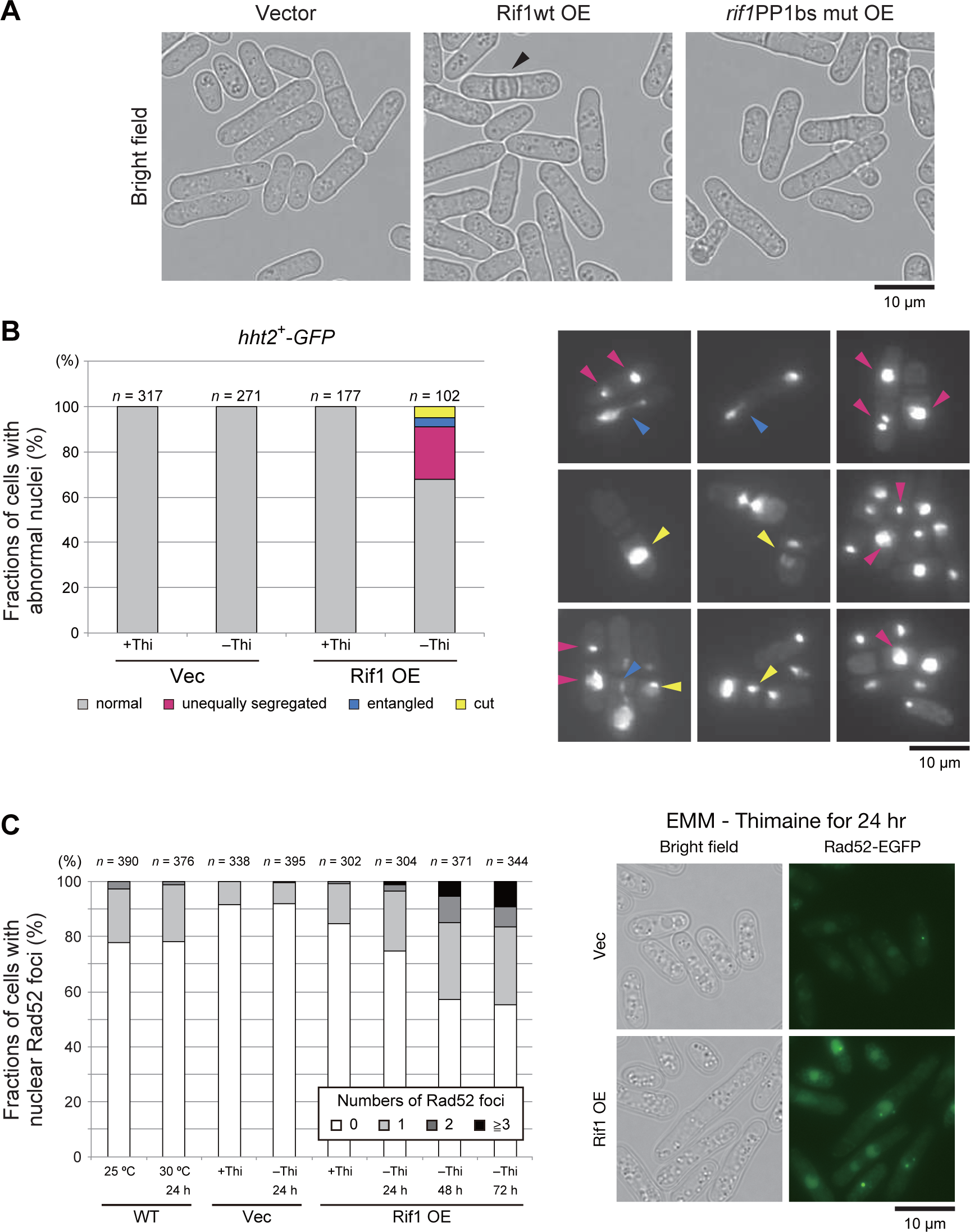
Rif1 overexpression induces unequal chromosome segregation and DNA damages. (**A**) Effect of Rif1 overexpression on morphology of cells observed under microscope. Arrowheads indicate cells with mitotic catastrophe ‘cut’ phenotype. (**B**) Chromosomes are visualized by *hht2* (histone H3 h3.2)-GFP (right) and the chromosome segregation was assessed in Rif1-overexpressing yeast cells. Cells with unequally segregated chromosomes (indicated by mazenta arrowheads) or entangled chromosomes (indicated by blue arrowheads) or cells showing cut phenotype (indicated by yellow arrowheads) increased at 24 hr after Rif1 overexpression (left). (**C**) Rif1 was overexpressed in cells expressing Rad52-EGFP by depletion of thiamine for 24, 48 and 72 hr. Rad52-EGFP foci in the cells were observed under fluorescent microscopy. The numbers of Rad52 foci (representing DNA damages) were counted, and cells containing 0, 1, 2 or >3 foci were quantified. The extent of DNA damages increased with the duration of Rif1 overexpression.

By using *GFP-α2tub* cells, we counted cells with spindle microtubules. In control cells (vector plasmid) and in the cells carrying pREP41-Rif1-Flag_3_ grown with thiamine, most cells showed only cytoplasmic microtubules and roughly only 1% of cells showed spindle microtubules; either short or long spindles were detected in roughly 0.5% each of the cell population (**Fig. 5A**). However, Rif1 overexpressing cells showed short spindles in up to 6% of the cell population and the population with long spindles decreased to one-half of the non-induced cells (**Fig. 5A**). This result unexpectedly suggested that at least 5-6% of cells overexpressing Rif1 arrest mitosis in metaphase-anaphase transition. The above results suggest a possibility that spindle assembly checkpoint is induced by Rif1 overexpression. Therefore, we examined the effect of *mad2* and *bub1* (required for spindle assembly checkpoint) mutation on the appearance of cells with spindles upon Rif1 overexpression (Bernard, Hardwick, & Javerzat, 1998; Bernard, Maure, & Javerzat, 2001; Garcia, Vardy, Koonrugsa, & Toda, 2001; Ikui, Furuya, Yanagida, & Matsumoto, 2002). The population of the cells with short spindle microtubules decreased to 1% or less in these mutants (**Fig. 5B**), indicating that the formation of short spindles depends on SAC. We therefore examined whether SAC is induced by overexpression of Rif1. When SAC is activated, APC/Cdc20 ubiquitin ligase is inhibited. This would stabilize Securin (Cut2) and CyclinB. We then measured the effects of Rif1 overexpression on the duration of Cut2 signal together with the locations of Sad1 (spindle pole body). In the control cells, the spindle appeared at 4 min and disappeared by 12 min. The Cut2 signal disappeared at around 14 min. In contrast, in Rif1-overproducing cells, the spindle appearing at time 0 was still visible at 36-38 min. The Cut2 signal persisted even after 30 min (**Fig. 5C**). These results show that SAC is activated by Rif1 overexpression. We then examined the effect of SAC mutations on the growth inhibition by Rif1-overexpression. Rif1 overexpression inhibited growth in *mad2*Δ and *bub1*Δ cells, indicating that growth inhibition is not caused by SAC (**Fig. 5D**). It is of interest that Rif1 overexpression inhibited growth more vigorously in the SAC mutants than in the wild-type cells. Indeed, after Rif1 overexpression, cells with cut phenotype increased from 4% in the wild-type cells to 8 % in *mad2*Δ and *bub1*Δ cells (**Fig. 5E**). These results suggest that SAC activation may partially suppress the cell death-inducing effect of Rif1 overexpression.

**Figure 5.**
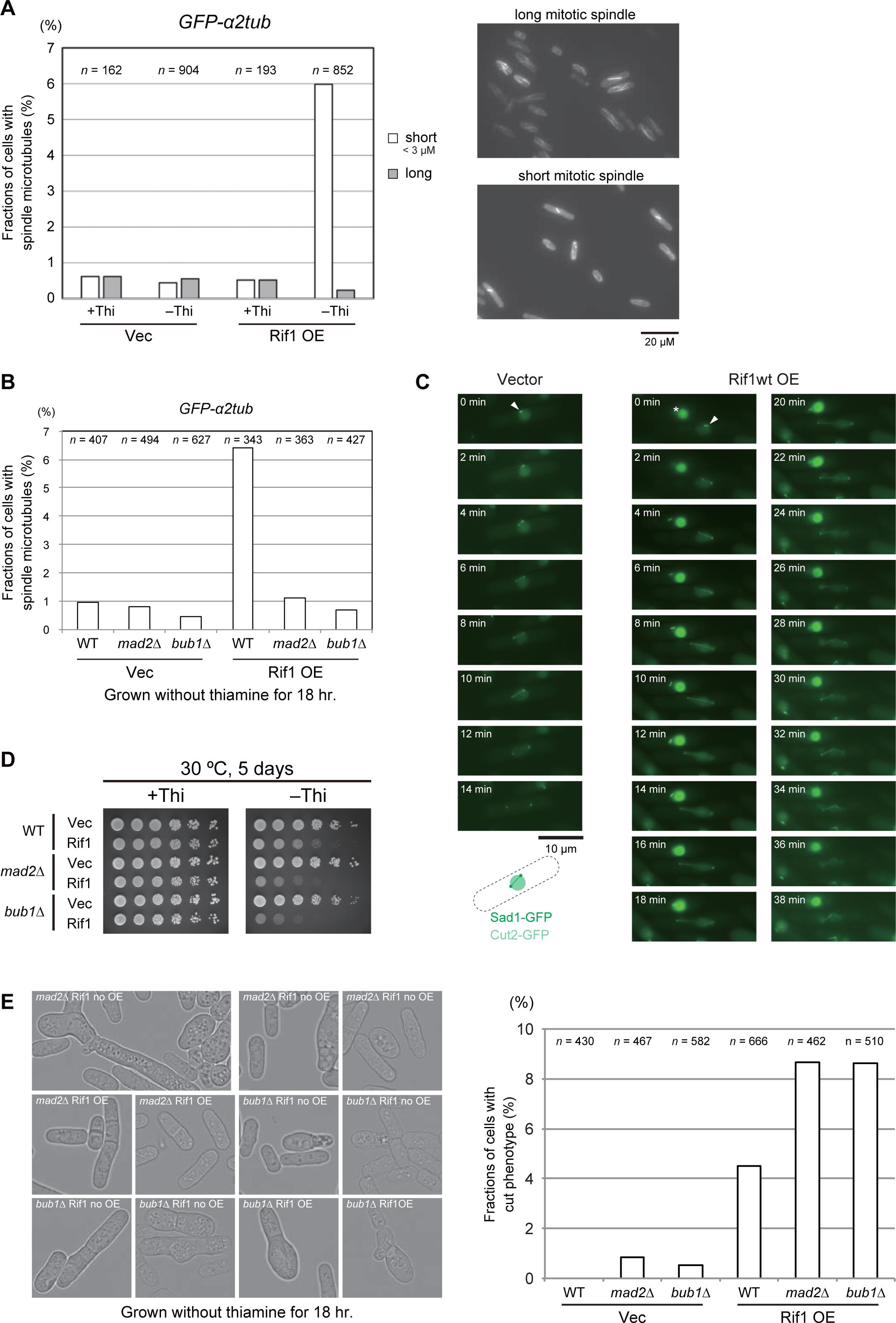
Cells with short tubulin spindles are accumulated in Rif1-overexpressing cells in a manner dependent on spindle assembly checkpoint (SAC) **(A)** Rif1 was overexpressed in cells expressing GFP-α2tub and cells with short or long spindle microtubules were counted. In the right panels, the photos of cells with short mitotic spindles and those with long spindles are shown. (**B**) Rif1 was overexpressed in the spindle assembly checkpoint activation mutants, *mad2*Δ or *bub1*Δ, and cells with spindle microtubules were counted. (**C**) SAC is induced in Rif1-overproducing cells. Cells expressing Sad1-GFP (spindle pole body) and Cut2-GFP (securin) were monitored under fluorescent microscope starting from the time when spindle pole bodies started to separate. Mitotic spindles between the two SPB disappear and the nuclear Cut2 signal disappear in non-overproducing cells by 14 min, while those in Rif1 overexpressing-cells stay as late as for 38 min. White arrowheads indicate Sad1. The drawing shows the nuclear signals of Cut2 (pale green) and two dots of Sad2 and connecting microtubules. The strong green signals indicating by * in Rif1wt OE samples represent a dead cell. (**D**) Spot tests of SAC mutant cells overexpressing Rif1. (**E**) Fractions of cells with cut phenotype are scored in the wild-type, *mad2*Δ or *bub1*Δ cells overproducing the wild-type Rif1. Left, phase contrast images of the cells; right, quantification of cut cell populations. Rif1 OE, Rif1 overexpression. In **A**, **B** and **E**, cells were grown in medium lacking thiamine for 18 hr.

The results indicate the aberrant chromatin segregation is responsible for growth inhibition and cell death. We noted that cells with aberrant spindles increased in cells overexpressing Rif1, and the fractions containing these structures were greater in PP1bs mutant overexpressing cells than in the wild-type Rif1 overexpressing cells (**Supplementary data Fig. S3**). In contrast, the populations of the cells with short spindles decreased with PP1bs mutant compared to the wilt-type Rif1. This is due to decreased level of SAC activation in the cells overexpressing PP1bs mutant than in those overexpressing the wild-type Rif1.

### Chromatin-binding of Rif1 is necessary for growth inhibition by Rif1 overexpression, and overexpressed Rif1 induces relocation of chromatin to nuclear periphery

We previously screened for *rif1* point mutants which suppress *hsk1-89*, and obtained two mutants, R236H and L848S, each of which alone could suppress *hsk1-89* (Kobayashi et al., 2019). R236H bound to Rif1bs (Rif1bs_I:2663_ and Rif1bs_II:4255_) and to a telomere as efficiently as the wild type in ChIP assays. On the other hand, L848S did not bind to either of the two Rif1bs or to the telomere (Kobayashi et al., 2019). We examined the effect of overexpression of these point mutants in wild-type and *hsk1-89* cells. R236H which can bind to chromatin caused growth defect in both wild type and *hsk1-89* when overexpressed (**Fig. 6A**). On the other hand, L848S which is compromised in chromatin binding activity showed no or very little growth inhibition in the wild type. Interestingly, L848S inhibited the growth of *hsk1-89* (**Fig. 6A**). Both mutant proteins were expressed at a level similar to that of the wild-type (data not shown).

**Figure 6.**
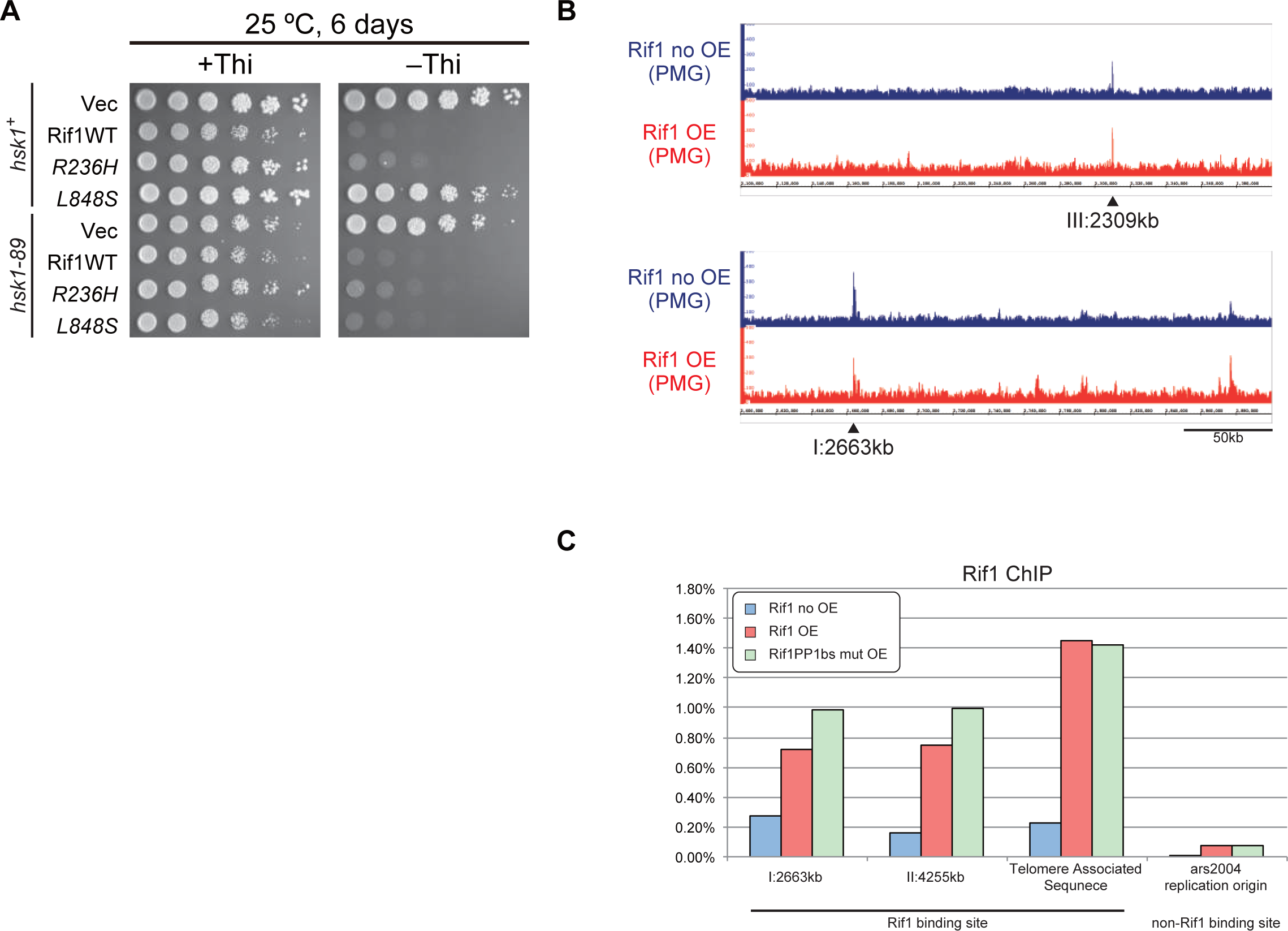
Requirement of chromatin binding for growth inhibition and chromatin binding profile of overexpressed Rif1. (A) Rif1 mutants were overexpressed in the wild-type and *hsk1-89* cells, and spot tests were conducted. R236H mutant binds to chromatin but L848S mutant does not (Kobayashi *et al*. 2019). (**B**) KYP1268 (*nda3-KM311*, Rif1-His_6_-Flag_10_; blue) and MS733 (*nda3-KM311*, nmt1-Rif1-His_6_-Flag_10_; red) were cultured in PMG medium containing 15 µM thiamine. The cells were washed with fresh PMG medium without thiamine and grown at 30°C for 12 hr. The cells were arrested at M-phase by shifting to 19.5°C for 6 hr, and then were released from M-phase by addition of an equal volume fresh PMG medium pre-warmed at 43°C. At 20 min after release, the cells were analyzed by ChIP-seq. Two known Rif1bs are indicated by arrowheads. (**C**) Binding of Rif1 to Rif1bs_I:2663kb_, Rif1bs_II:4255kb_, TAS (telomere of chromosome II) and ars2004 (non-Rif1bs) was measured in the wild-type cells harboring vector, pREP41-Rif1-Flag_3_, or pREP41-*rif1*PP1bsmut-Flag_3_ by ChIP-qPCR. Cells were grown in the medium lacking thiamine for 18 hr before harvest. The IP efficiency was normalized by the level of input DNA.

The above results strongly suggest that chromatin binding of Rif1 is important for growth inhibition. Therefore, we have examined chromatin binding of overexpressed Rif1 protein by ChIP seq analyses. The results indicate that overexpressed Rif1 binds to multiple sites on the chromatin, in addition to its targets in the non-overproducing wild-type cells (**Fig. 6B**). ChIP-qPCR showed that overexpressed Rif1 binds to known Rif1bs sequences as well as to a telomere with 3 to 7 fold higher efficiency than the endogenous Rif1 does, and binds also to a non-Rif1bs sequence (**Fig. 6C**). There results suggest a possibility that the aberrant chromatin binding of Rif1 may be related to the induction of aberrant chromatin morphology and resulting growth inhibition and cell death.

We examined the chromatin morphology by using the cells containing GFP-labeled histone (h3.2-GFP). Interestingly, induction of Rif1 expression led to increased cell populations carrying nuclei with chromatin enriched at the nuclear periphery. This population reached over 6% with the wild type and 11 % with the PP1bs mutant at 18 hr after induction (**Fig. 7A and 7B**). Enrichment of chromatin at nuclear periphery could be caused by enlarged nucleoli as a result of Rif1 overproduction. We therefore measured the sizes of nucleoli by labeling Gar2 protein. We did not detect any effect on the sizes of nucleoli by overexpression of the wild-type or PP1bs mutant Rif1 protein (**Supplementary Fig. S4**), showing that chromatin relocation is not caused by enlarged nucleoli. The higher level of aberrant chromatin caused by the PP1bs mutant may be consistent with more severe growth inhibition with this mutant. The chromatin binding-deficient L848S mutant did not significantly induce relocation of chromatin, whereas R236H mutant, that is capable of chromatin binding, induced the relocation in 5% of the population (**Fig. 7C and D**), in keeping with growth-inhibiting properties of the latter mutant but not of the former.

**Figure 7.**
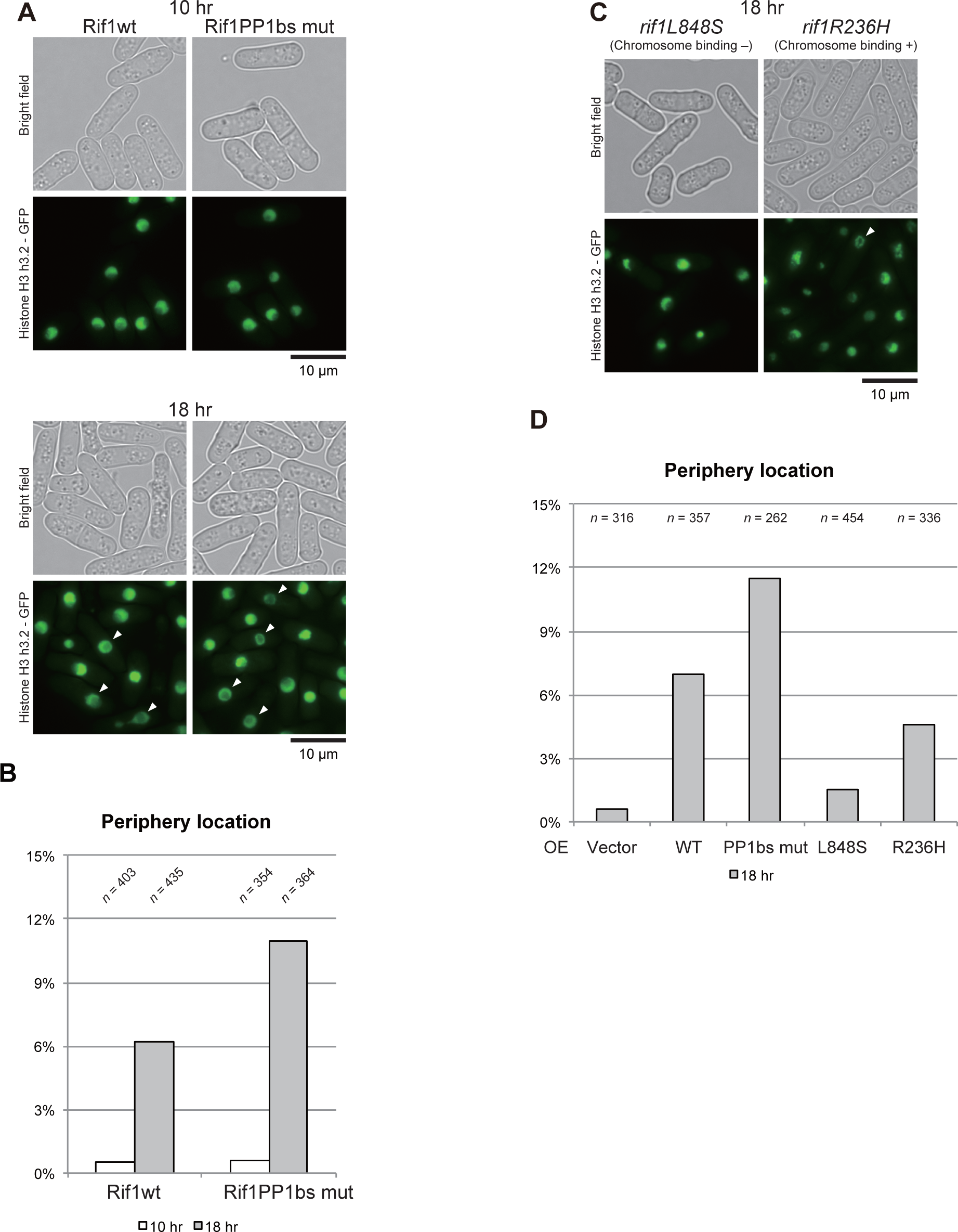
Chromatin morphology of the cells after induction of Rif1 expression. Cells expressing GFP-fused Histone H3 (h3.2-GFP) were observed under fluorescent microscope after induction of Rif1 protein (wild-type or PP1bs mutant in **A** and L848S [chromatin binding-deficient] and R236H [chromatin binding-proficient] mutants in **C**) for 10 hr (upper panel of **A**) and 18 hr (lower panel of **A,** and **C**). Phase contrast and fluorescent images of the cells (**A** and **C**) and quantification of cells with chromatin relocated at the nuclear periphery (**B** and **D**) are presented. Cells with chromatin relocated at the nuclear periphery are indicated by arrowheads.

### Nuclear dynamics of Rif1 protein

In order to visualize nuclear dynamics of Rif1 protein, we have fused a fluorescent protein to Rif1. Rif1 protein in higher eukaryotes contains a long IDP (intrinsically disordered polypeptide) segment between the N-terminal HEAT repeat sequences and C-terminal segment containing G4 binding and oligomerization activities. The fission yeast Rif1 does not carry IDP, but contains HEAT repeats and the C-terminal segment with similar biochemical activities. Thus, we speculated that insertion of a foreign polypeptide at the boundary of the two domains would least affect the overall structure of the protein, and introduced the mKO2 DNA fragment at aa1090/1091 or at aa1128/1129 (**Supplementary Fig. S5 A**). The resulting plasmid DNAs were integrated at the endogenous *rif1* locus generating MIC11-130 *rif1*-mKO2-1 and MIC12-123 *rif1*-mKO2-2, respectively.

We then evaluated the functions of the fusion proteins. *hsk1-83* (ts) cells did not grow at 30°C (non-permissive temperature), whereas *hsk1-83 rif1*Δ cells did. On the other hand, *hsk1-83 rif1*-mKO2 did not grow at 30°C (**Supplementary Fig. S5 B**), indicating that the Rif1-mKO2 retains the wild-type replication-inhibitory functions. In order to evaluate their telomere functions, we examined the telomere length in Rif1-mKO2 cells. As reported, the telomere length increased in *rif1*Δ cells (**Supplementary Fig. S5 C**, lane 2), whereas that in Rif1-mKO2-1 and -2 cells did not significantly change (**Supplementary Fig. S5 C**, lanes 3 and 4), suggesting that the insertion of mKO2 does not affect the Rif1 function at telomeres. We chose Rif1-mKO2 (aa1090/1091) carrying cells for further analysis. Rif1-mKO2 exhibited strong dots in nuclei (**Supplementary Fig. S5 D and E**), which coloclalized with Taz1-GFP or Rap1-EGFP (**Supplementary Fig. S5 D**, data not shown), indicating that they represent telomeres. Minute foci appeared in nuclei, probably representing Rif1 bound to the chromosome arms. Thus, Rif1-mKO2 (aa1090/1091) cells permit the visualization of dynamics of the endogenous Rif1 protein. Time laps analyses of Rif1-mKO2 revealed a few big foci in each cell which colocalize with Taz1 along with minute other nuclear foci that are highly dynamic and represent Rif1 on chromatin arms (**Supplementary movies; Supplementary Fig. S5 D**).

### Overexpression of Rif1 causes relocation of the endogenous Rif1 protein

Upon overexpression of Rif1, either wild-type or a PP1bs mutant, in Rif1-mKO2 cells, mKO2 signals spread through nuclei (**Supplementary Fig. S5 E**), consistent with the promiscuous chromatin binding of overexpressed Rif1. The overexpressed Rif1 would form mixed oligomers with the endogenous Rif1-mKO2, relocating some of the telomere-bound Rif1-mKO2 to chromosome arms. Prewash with 0.1% Triton X-100 and DNase I before PFA fixation led to appearance of multiple clear dots in nuclei in Rif1-mKO2 cells (**Fig. 8A** and **Supplementary Fig. S6 A**), since at least some of the Rif1 bound to chromatin arms is resistant to Triton/DNaseI pretreatment and that on telomere is more sensitive. Overexpression of Rif1 resulted in increased numbers of dots (**Fig. 8A, B** and **Supplementary Fig. S6 A, B, C**), consistent with the relocation of Rif1-mKO2 from the telomere to nuclear matrix-related insoluble compartments. The overall nuclear fluorescent intensities of the prewashed mKO2-Rif1 cells also increased after Rif1 overexpression compared to the vector control (**Fig. 8C** and **Supplementary Fig. S6 C**), consistent with above speculation. These results support the conclusion that overexpressed Rif1 promotes aberrant tethering of chromatin to nuclear membrane/ nuclear matrix-related insoluble compartments and that the resulting aberrant chromatin organization causes mitotic defect.

**Figure 8.**
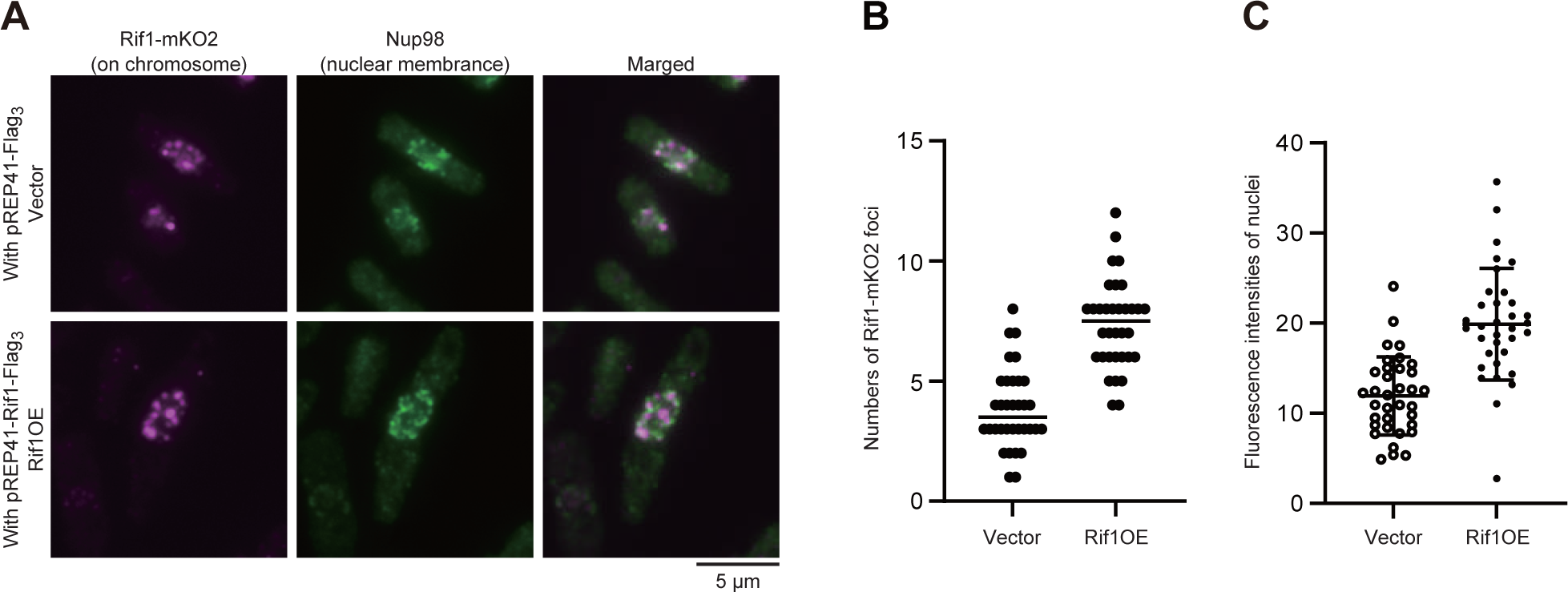
The endogenous Rif1 protein was relocated upon overexpression of Rif1. Rif1-mKO2 cells, in which the endogenous Rif1 was tagged with mKO2, were grown in the absence of thiamine for 20 hr, and were extracted by Triton X-100 and DNase I and remaining endogenous Rif1-mKO2 signals (mazenta) were observed. The nuclear envelope was stained with Nup98 antibody (green). The numbers (**B**) and the intensities (**C**) of nuclear foci were quantified in Rif1-mKO2 cells harboring pREP41-Flag_3_ (Vector) or pREP41-Rif1-Flag_3_ (Rif1OE) grown as in (**A**).

## Discussion

Rif1 is an evolutionary conserved nuclear factor that plays roles in various chromosome transactions including DSB repair, DNA replication, transcription, and epigenetic regulation. Rif1 proteins from fission yeast and mammalian cells bind to G4 structure, and generate higher-order chromatin architecture (Y. Kanoh et al., 2015; Moriyama, Yoshizawa-Sugata, & Masai, 2018). Both fission yeast and human Rif1 are biochemically enriched in nuclear insoluble fractions, and a portion of mammalian Rif1 is localized at nuclear periphery. It was speculated that Rif1 tethers chromatin fiber along the nuclear membrane, generating a chromatin compartment in the vicinity of nuclear periphery (Kobayashi et al., 2019; Yamazaki et al., 2012). However, effects of Rif1 on chromatin localization in nuclei and its subsequent outcome have not been explored.

The fission yeast genome contains 35 significant Rif1 binding sites (Rif1bs), as determined by ChIP-seq analyses, and Rif1 binding to these G4-forming Rif1bs generates DNA regions spanning the binding sites as long as ∼100 kb in which origin firing is suppressed (Y. Kanoh et al., 2015). In contrast, mammalian Rif1 binds extensively to late replicating segments, concomitant with occasional strong binding to sequences containing multiple G-tracts and capable of forming G4 (Foti et al., 2016; Klein et al., 2021)(Yoshizawa et al, unpublished data). The copy number of Rif1 in fission yeast was estimated to be ∼2,000 molecules/ cell (Masai et al., unpublished data).

Previous biochemical analyses indicate that the C-terminal segment may form oligomers (8-mer to 16-mer). Thirty-five binding sites account for ∼500 molecules of Rif1. The remaining Rif1 molecules may be bound to telomeres. Increase of the Rif1 molecule numbers may influence the binding profile of Rif1, and may affect S phase or cell cycle progression through alteration of chromatin architecture.

In this communication, we report that overexpressed Rif1 protein inhibits growth and induces cell death in fission yeast cells. This effect is independent of its ability to recruit PP1. Rif1 overexpression induces S phase delay in a manner dependent on PP1 binding. Replication checkpoint is activated by the PP1bs mutant but not by the wild-type Rif1. SAC is also activated and cells with short spindle accumulate notably by the overexpression of the wild-type Rif1 but not by that of the PP1bs mutant or in PP1 mutant cells (data not shown). The primary cause for the cell death appears to be defective mitosis, as indicated by the appearance of aberrant microtubule spindles. The overexpressed Rif1 binds to chromatin at sites that are not normally bound in the wild-type cells, and nuclear DNAs are tethered to the nuclear periphery, consistent with the potential role of Rif1 in tethering chromatin fiber at the nuclear membranes. Our findings also suggest potential novel roles of Rif1 in regulation of mitosis, notably the proper function of microtubule spindles.

### Events induced by overexpression of Rif1 in fission yeast cells

We first observed severe growth inhibition of fission yeast cells by overexpression of Rif1 protein. Rif1 overexpression delayed S phase initiation and progression in synchronized cell populations. Interestingly, S phase inhibition was not observed by overexpression of the PP1bs mutant of Rif1, indicating that inhibition of DNA synthesis depends on the recruitment of PP1 (**Fig. 3A**). Replication checkpoint, as measured by Cds1 kinase activity, was activated by overexpression of the PP1bs mutant of Rif1 protein, but not by the wild-type Rif1 protein (**Fig. 3C**). This could be due to the ongoing S phase in PP1bs mutant-overexpressing cells while S phase is inhibited by the wild-type Rif1. Ongoing replication forks would be interfered by the bound Rif1 proteins, activating replication checkpoint. On the other hand, in the wild-type Rif1 overexpression, initiation of DNA synthesis is blocked by Rif1, thus generating less fork blocks, i.e. less replication checkpoint activation.

Cells with short spindles accumulate in Rif1-overproduced cells, and this incident depends on SAC (**Fig. 5B**). Growth inhibition is observed in SAC mutant cells, indicating that short spindle formation is not critical for growth inhibition. Rather, severer growth inhibition was observed in the SAC mutants. Thus, SAC activation may be protective against Rif1-induced growth inhibition and cell death. Short spindles were reported to accumulate in SAC activated cells, and it was reported that replication intermediates or recombination-intermediates can activate SAC (Nakano et al., 2014), resulting in cells with short spindles. SAC was less strongly activated by overexpression of a PP1bs mutant, consistent with reduced short spindles in this mutant. Rif1 inhibited growth more strongly in SAC mutants than in the wild-type cells. The PP1bs mutant inhibited growth slightly more strongly than the wild-type Rif1. These results indicate that SAC-induced mitotic arrest with short spindles may serve as a protective barrier against aberrant mitosis that would lead to cell death (**Fig. 9**).

**Figure 9.**
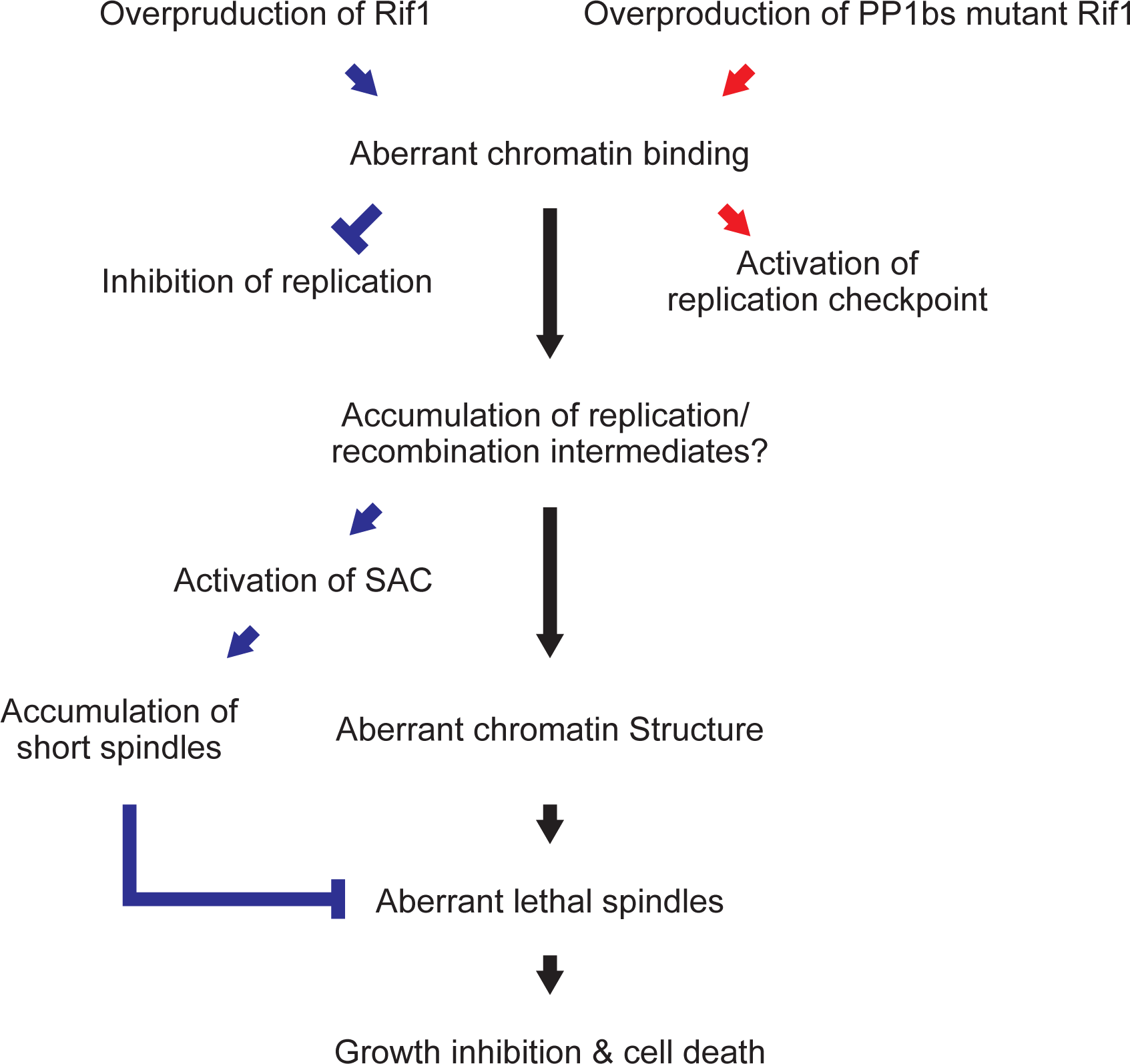
Cellular events induced by overexpression of Rif1 in fission yeast. Overproduction of Rif1 leads to its aberrant chromatin binding and inhibits S phase initiation and progression through its ability to recruit PPase. Excessive chromatin binding of Rif1 results in aberrant tethering of chromatin fibers to nuclear periphery, which may directly or indirectly inhibit proper progression of chromosome segregation, eventually leading to cell death. Overexpression of the wild-type Rif1 inhibits DNA replication, whereas that of PP1bs mutant Rif1 does not inhibit DNA replication but activates replication checkpoint. Rif1 overexpression induces SAC, leading to increased cell population with short spindles, which probably antagonizes induction of aberrant chromosome structures.

### Aberrant chromatin structures induced by Rif1 suggests the ability of Rif1 to tether chromatin at nuclear periphery

Histone H3 (H3.2) labeled chromatin is detected uniformly in nuclei in the wild-type cells. Upon overexpression of Rif1 protein, nuclei with chromatin enriched at the nuclear periphery were observed in more than 6% or 10% of the cells with the wild-type or PP1bs mutant Rif1, respectively. Furthermore, chromatin binding-deficient mutant, L848S, failed to induce chromatin relocation to nuclear periphery (**Fig. 7**). Our results are consistent with the idea that Rif1 promotes association of chromatin with nuclear membrane through its chromatin binding ability and through its potential membrane association ability. Fission yeast Rif is present in highly insoluble fractions in biochemical fractionation (Y. Kanoh et al., 2015), and mammalian Rif1 is localized at nuclear periphery during G1-S phase (Yamazaki et al., 2012). Budding yeast Rif1 is palmitoylated and anchored to membrane (Fontana et al., 2019). These results suggest that fission yeast Rif1 protein is also tethered to the nuclear membrane. The density map prediction indicated the enrichment of Rif1 at nuclear periphery and around nucleoli, where DNA replication occurs predominantly in the absence of Rif1 (**data not shown**). Overexpressed Rif1 binds to genomic DNA more promiscuously and the Rif1-bound chromatin fiber would be tethered along the nuclear membrane/ detergent- and DNase I-insoluble compartments in the presence of overexpressed Rif1. Thus, overexpression of Rif1 would lead to relocation of the chromatin segment located in the interior of nuclei (early-replicating loci) to nuclear periphery. Replication of DNA in the vicinity of chromatin tethered segment would be inhibited upon recruitment of PP1. It was previously reported that artificial tethering of an early-firing origin at nuclear periphery did not render it late-firing in budding yeast (Ebrahimi et al., 2010). This is consistent with our result that Rif1-mediated chromatin recruitment at nuclear membrane alone does not inhibit S phase, and the recruitment of PP1 by Rif1 is essential for the inhibition.

### Cellular dynamics of Rif1 protein

By using a Rif1 derivative containing mKO2 between the HEAT repeat and C-terminal segment, cellular Rif1 dynamics was examined. In addition to the strong dots corresponding to telomeres (**Supplementary Fig. S5 D and E**), fine dots representing arm binding were observed ((Klein et al., 2021)**Supplementary movie**). Upon overexpression of Rif1, the endogenous Rif1-derived mKO2 signals relocated from telomeres to the entire areas of nuclei (**Supplementary Fig. S5 E**), indicating that overexpressed Rif1, forming multimers with Rif1-mKO2, spreads over the chromosome arms. The prewash of nuclei with detergent and DNase I enhanced the Rif1 signals in nuclei, notably at nuclear periphery (**Supplementary Fig. S6 B**). Both the numbers of foci and overall intensities of nuclear signals increased upon Rif1 overproduction (**Fig. 8B and C**), presumably due to relocation of endogenous Rif1 from telomere to chromosome arms and to detergent- and DNase I-resistant insoluble compartments through mixed oligomer formation with the overexpressed Rif1.

### Mechanisms of formation of aberrant microtubule spindles

Rif1 overexpression ultimately induces cell death through aberrant mitosis. In addition to cells with short spindles, those with aberrant defective microtubules appear. The fractions of these cells increase in PP1 mutant cells as well as in PP1bs mutant-overproducing cells. PP1-Rif1 interaction is regulated by phosphorylation of Rif1 and Aurora B kinase was reported to play a major role in phosphorylating the PP1bs, promoting the dissociation of PP1 from Rif1 during M phase (Bhowmick et al., 2019; Nasa, Rusin, Kettenbach, & Moorhead, 2018). Overexpressed Rif1 would recruit PP1, counteracting the phosphorylation events by Aurora B and other kinases essential for mitosis. This may lead to misconduct in mitotic events. However, since PP1bs mutants can also cause aberrant microtubules and cell death, PP1 may not be the primary cause for aberrant microtubule cell death. The aberrant chromatin structure caused by overexpressed Rif1 may affect the mitotic chromatin structure, leading to mitotic defect. Alternatively, the aberrant association of Rif1 with mitotic kinases and potentially with microtubules could directly be linked to deficient microtubules in Rif1-overproducing cells.

## Data Availability

The reagents, Oligos, plasmids and Strains in this study are listed in tables 1∼4.

## Acknowledgements

We thank Rino Fukatsu and Naoko Kakusho for excellent technical assistance. We thank Justin O’Sullivan for providing us with the data on prediction of nuclear localization of Rif1 and DNA replication in fission yeast cells. This paper is dedicated to Dr. Seiji Matsumoto, a dearest friend and collaborator, who passed away on November 22, 2020, after a long fight against pancreatic cancer.

## Material and Methods

### Medium for Schizosaccharomyces pombe

YES media contains 0.5 % yeast extract (Gibco), 3 % glucose (FUJIFILM Wako) and 0.1 mg/mL each of adenine (Sigma-Aldrich), uracil (Sigma -Aldrich), leucine (FUJIFILM Wako), lysine (FUJIFILM Wako) and histidine (FUJIFILM Wako). YES plates were made by adding 2 % agar (Gibco) to YES media. Synthetic dextrose minimal medium (SD) contains 6.3 g/L Yeast Nitrogen Base w/o Amino Acids (BD), 2% glucose and 0.1 mg/mL each of required amino acids. Edinburgh Minimal Medium (EMM) contains 12.3 g/L EMM BROTH WITHOUT NITROGEN (FORMEIUM), 2% glucose and 0.1 mg/mL each of required amino acids. Pombe Minimal Glutamate (PMG) contains 27.3 g/L EMM BROTH WITHOUT NITROGEN, 5 g/L L-glutamic acid (Sigma-Aldrich) and 0.1 mg/mL each of required amino acids. Fifteen µM thiamine (Sigma-Aldrich) was added to EMM or PMG medium to repress the nmt1 promoter activity. Yeast strains and plasmids used in this study are listed in **Table 1**.

**Table 1.** Strains list used in this study.

| **Strain** | **Genotype** | **Source** |
| --- | --- | --- |
| YM71 | h– leu1-32 ura4-D18 | Our Stock |
| MS104 | h– leu1-32 ura4-D18 hsk1-89:ura4+ | Our Stock |
| KYP1827 | h– leu1-32 ura4-D18 Rif1-Flag3:kanR pREP41 | Our Stock |
| JX503 | h– dis2-11 leu1 | Our Stock |
| JX504 | h– dis2::ura4+ leu1 ura4 | Our Stock |
| FY9620 | h– leu1 ura4 sds21::ura4+ | NBRP |
| MS672 | h– leu1-32 ura4-D18 rif1Δ::ura4+ | Our Stock |
| KYP1268 | h– nda3-KM311 leu1-32 ura4-D18 Rif1-6xHis-10xFlag | Our Stock |
| MS733 | h– nda3-KM311 leu1-32 ura4-D18 rif1::Pnmt-rif1-6xHis-10xFlag:kan | Our Stock |
| KYP1283 | h– nda3-KM311 leu1-32 ura4-D18 Pnmt1-rif1PP1BSmut-6xHis-10xFlag:kanR | Our Stock |
| FY15623 | h– ade6-M216 leu1-32 ura4-D18 hht2+-GFP::ura4+ | NBRP |
| MS360 | h– leu1-32 ura4-D18 rad22-YFP:KanR | Our Stock |
| KYP011 | h– leu1-32 ura4-D18 Rad52-EGFP:kanR | Our Stock |
| MS130 | h+ leu1-32 ura4-D18 lys1+::pmt1-GFP-alpha2tub | Our Stock |
| KYP1801 | h– leu1-32 ura4-D18 mad2::ura4+ (AE147) lys1+::pmt1-GFP-alpha2tub | Our Stock |
| KYP1802 | h– bub1::ura4+ leu1 ade6-M216 lys1+::pmt1-GFP-alpha2tub | Our Stock |
| 23-B10 | h90 ade6-M216 leu1 his3-D1 cut2-GFP<<kanR sad1-GFP<<kanR | Gifted from Dr. Ueno |
| FY14161 | h– leu1-32 ura4-D18 taz1::ura4+ | NBRP |
| MS129 | h– leu1-32 ura4-D18 tel1-D1::kanMX4 rad3::ura4+ | Our Stock |
| NI485 | h+ leu1-32 ura4-D18 ade6 rad3::ura4+ | Our Stock |
| MS221 | h+ leu1-32 ade6-M216 ura4-D18 chk1::ura4+ | Our Stock |
| NI453 | h– leu1-32 ura4-D18 cds1::ura4+ | Our Stock |
| FY7826 | h– leu1-32 ura4-D18 mad2::ura4+ (AE147) | NBRP |
| MS182 | h– leu1-32 ura4-D18 cdc25-22 | Our Stock |
| MS285 | h– ade6-M210 Leu2-32 ura4-D18 rif1::LEU2 cdc25-22 | Our Stock |
| MIC2-11 | h– Leu2-32 ura4-D18 rif1:mKO2 | Our Stock |
| MIC11-130 | h– Leu2-32 ura4-D18 rif1:mKO2(1090) cdc25-22 | Our Stock |
| MIC12-123 | h+ Leu2-32 ura4-D18 rif1:mKO2(1128) cdc25-22 | Our Stock |
| NI324 | h+ ade6-M210 Leu2-32 ura4-D18 hsk1-89:ura4+ | Our Stock |
| MS744 | h– Leu2-32 ura4-D18 rif1:mKO2 hsk1-89:ura4+ | Our Stock |
| MS745 | h+ Leu2-32 ura4-D18 rif1:mKO2 hsk1-89:ura4+ | Our Stock |
| MS558 | h– Leu2-32 ura4-D18 hsk1-89:ura4+ rif1::ura4+ | Our Stock |
| MS195 | h– ade6-M216 leu1-32 ura4-D18 wee1-50 | Our Stock |

**Table 2.**
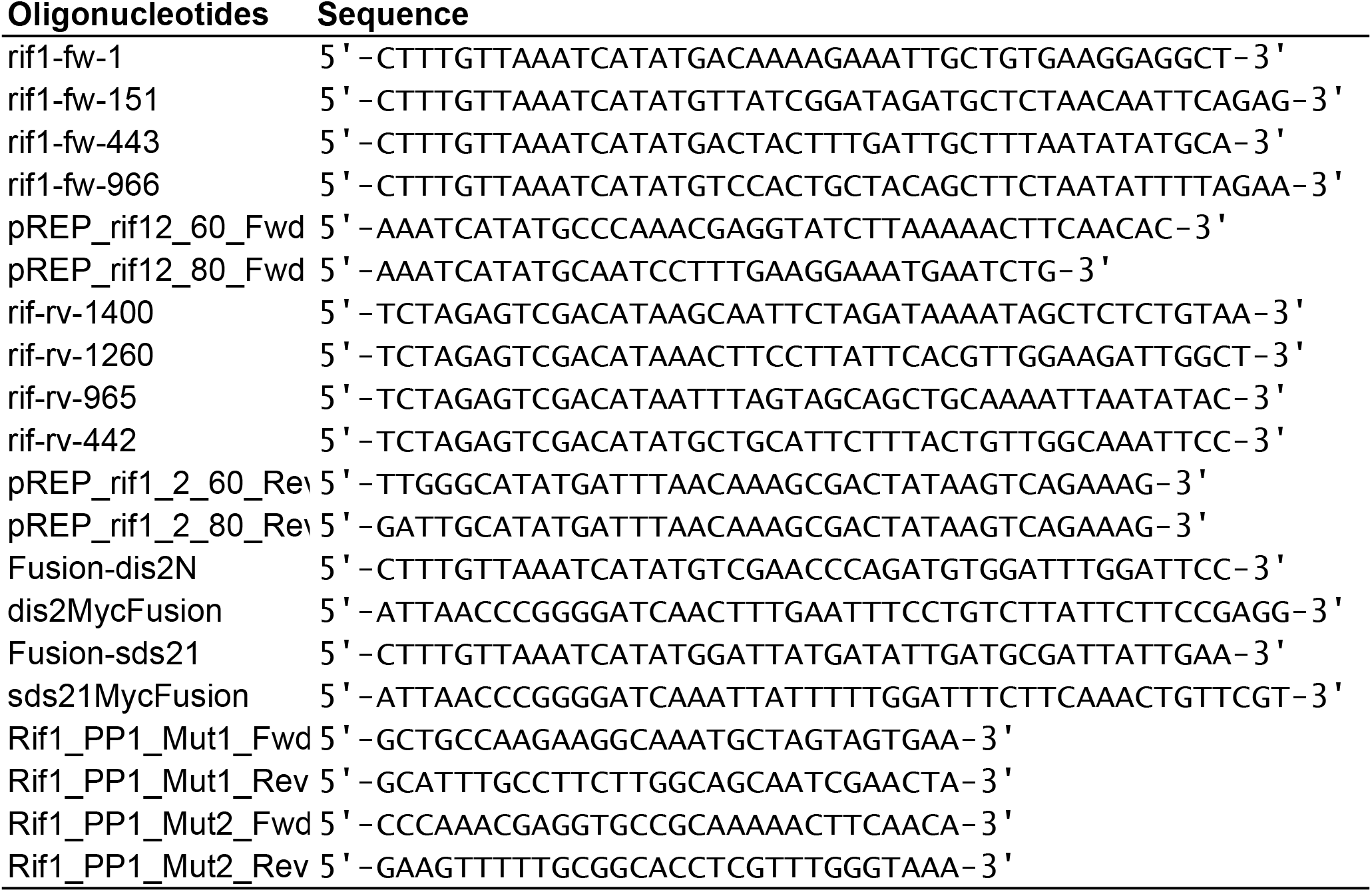

**Table 3.**
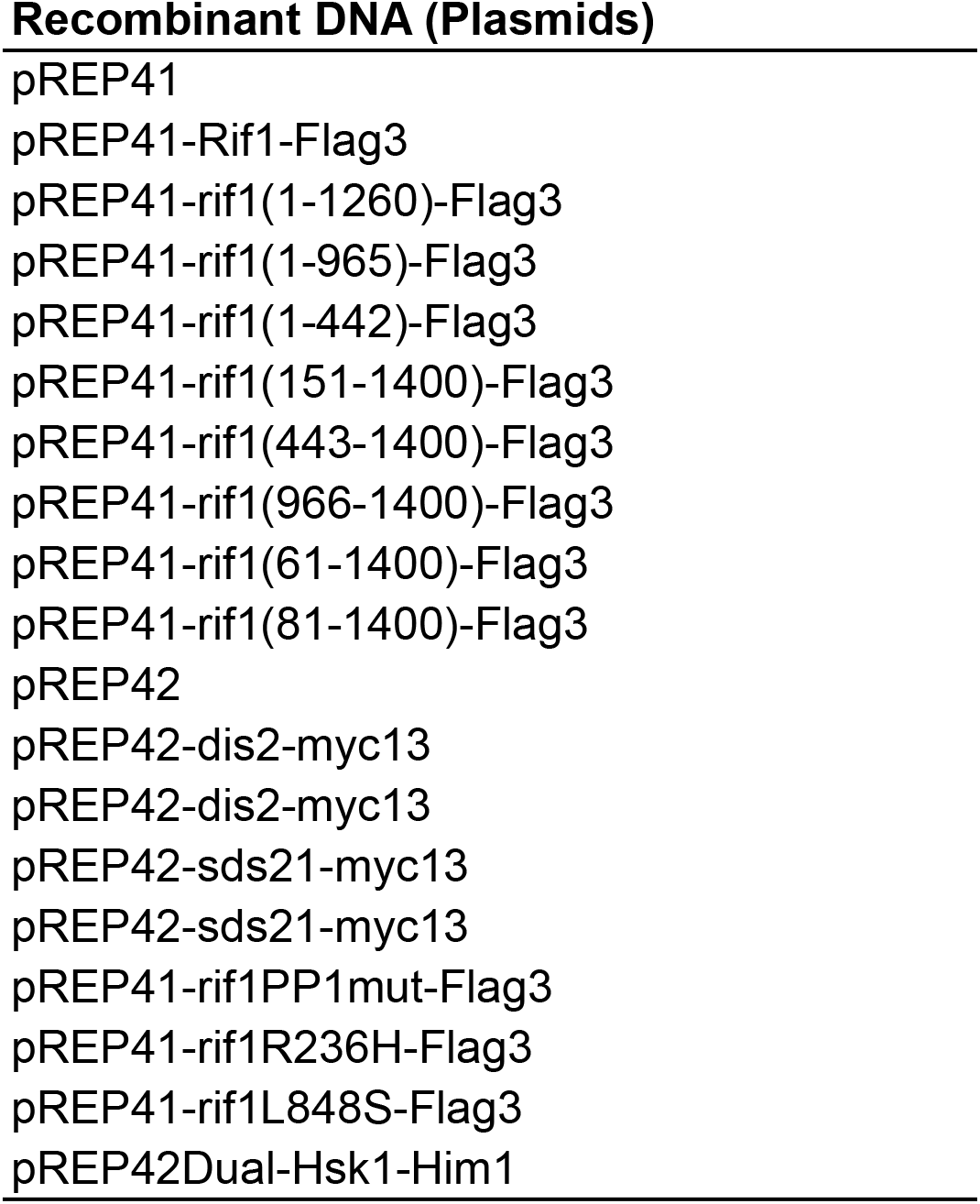

**Table 4.**
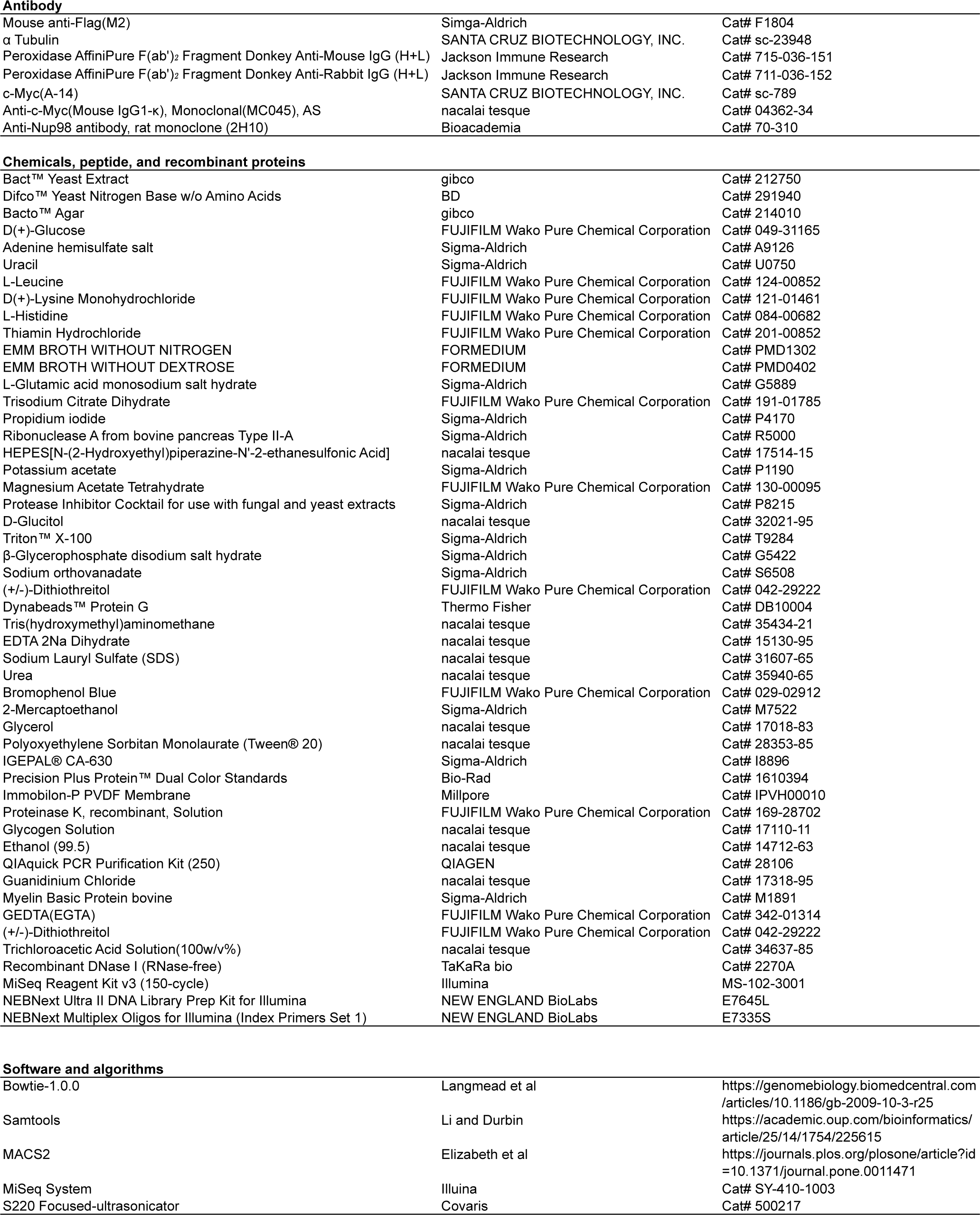
REAGENT or RESOURECE.

Antibody
|  |  |  |
| --- | --- | --- |
| Mouse anti-Flag(M2) | Simga-Aldrich | Cat# F1804 |
| $\alpha$ Tubulin | SANTA CRUZ BIOTECHNOLOGY, INC. | Cat# sc-23948 |
| Peroxidase AffiniPure F(ab')<sub>2</sub> Fragment Donkey Anti-Mouse IgG (H+L) | Jackson Immune Research | Cat# 715-036-151 |
| Peroxidase AffiniPure F(ab')<sub>2</sub> Fragment Donkey Anti-Rabbit IgG (H+L) | Jackson Immune Research | Cat# 711-036-152 |
| c-Myc(A-14) | SANTA CRUZ BIOTECHNOLOGY, INC. | Cat# sc-789 |
| Anti-c-Myc(Mouse IgG1-k), Monoclonal(MC045), AS | nacalai tesque | Cat# 04362-34 |
| Anti-Nup98 antibody, rat monoclonal (2H10) | Bioacademia | Cat# 70-310 |
| Chemicals, peptide, and recombinant proteins |  |  |
| Bact™ Yeast Extract | gibco | Cat# 212750 |
| Difco™ Yeast Nitrogen Base w/o Amino Acids | BD | Cat# 291940 |
| Bacto™ Agar | gibco | Cat# 214010 |
| D(+)-Glucose | FUJIFILM Wako Pure Chemical Corporation | Cat# 049-31165 |
| Adenine hemisulfate salt | Sigma-Aldrich | Cat# A9126 |
| Uracil | Sigma-Aldrich | Cat# U0750 |
| L-Leucine | FUJIFILM Wako Pure Chemical Corporation | Cat# 124-00852 |
| D(+)-Lysine Monohydrochloride | FUJIFILM Wako Pure Chemical Corporation | Cat# 121-01461 |
| L-Histidine | FUJIFILM Wako Pure Chemical Corporation | Cat# 084-00682 |
| Thiamin Hydrochloride | FUJIFILM Wako Pure Chemical Corporation | Cat# 201-00852 |
| EMM BROTH WITHOUT NITROGEN | FORMEDIUM | Cat# PMD1302 |
| EMM BROTH WITHOUT DEXTROSE | FORMEDIUM | Cat# PMD0402 |
| L-Glutamic acid monosodium salt hydrate | Sigma-Aldrich | Cat# G5889 |
| Trisodium Citrate Dihydrate | FUJIFILM Wako Pure Chemical Corporation | Cat# 191-01785 |
| Propidium iodide | Sigma-Aldrich | Cat# P4170 |
| Ribonuclease A from bovine pancreas Type II-A | Sigma-Aldrich | Cat# R5000 |
| HEPES[N-(2-Hydroxyethyl)piperazine-N'-2-ethanesulfonic Acid] | nacalai tesque | Cat# 17514-15 |
| Potassium acetate | Sigma-Aldrich | Cat# P1190 |
| Magnesium Acetate Tetrahydrate | FUJIFILM Wako Pure Chemical Corporation | Cat# 130-00095 |
| Protease Inhibitor Cocktail for use with fungal and yeast extracts | Sigma-Aldrich | Cat# P8215 |
| D-Glucitol | nacalai tesque | Cat# 32021-95 |
| Triton™ X-100 | Sigma-Aldrich | Cat# T9284 |
| β-Glycerophosphate disodium salt hydrate | Sigma-Aldrich | Cat# G5422 |
| Sodium orthovanadate | Sigma-Aldrich | Cat# S6508 |
| (+/-)-Dithiothreitol | FUJIFILM Wako Pure Chemical Corporation | Cat# 042-29222 |
| Dynabeads™ Protein G | Thermo Fisher | Cat# DB10004 |
| Tris(hydroxymethyl)aminomethane | nacalai tesque | Cat# 35434-21 |
| EDTA 2Na Dihydrate | nacalai tesque | Cat# 15130-95 |
| Sodium Lauryl Sulfate (SDS) | nacalai tesque | Cat# 31607-65 |
| Urea | nacalai tesque | Cat# 35940-65 |
| Bromophenol Blue | FUJIFILM Wako Pure Chemical Corporation | Cat# 029-02912 |
| 2-Mercaptoethanol | Sigma-Aldrich | Cat# M7522 |
| Glycerol | nacalai tesque | Cat# 17018-83 |
| Polyoxyethylene Sorbitan Monolaurate (Tween® 20) | nacalai tesque | Cat# 28353-85 |
| IGEPAL® CA-630 | Sigma-Aldrich | Cat# I8896 |
| Precision Plus Protein™ Dual Color Standards | Bio-Rad | Cat# 1610394 |
| Immobilon-P PVDF Membrane | Millipore | Cat# IPVH00010 |
| Proteinase K, recombinant, Solution | FUJIFILM Wako Pure Chemical Corporation | Cat# 169-28702 |
| Glycogen Solution | nacalai tesque | Cat# 17110-11 |
| Ethanol (99.5) | nacalai tesque | Cat# 14712-63 |
| QIAquick PCR Purification Kit (250) | QIAGEN | Cat# 28106 |
| Guanidinium Chloride | nacalai tesque | Cat# 17318-95 |
| Myelin Basic Protein bovine | Sigma-Aldrich | Cat# M1891 |
| GEDTA(EGTA) | FUJIFILM Wako Pure Chemical Corporation | Cat# 342-01314 |
| (+/-)-Dithiothreitol | FUJIFILM Wako Pure Chemical Corporation | Cat# 042-29222 |
| Trichloroacetic Acid Solution(100w/v%) | nacalai tesque | Cat# 34637-85 |
| Recombinant DNase I (RNase-free) | TaKaRa bio | Cat# 2270A |
| MiSeq Reagent Kit v3 (150-cycle) | Illumina | MS-102-3001 |
| NEBNext Ultra II DNA Library Prep Kit for Illumina | NEW ENGLAND BioLabs | E7645L |
| NEBNext Multiplex Oligos for Illumina (Index Primers Set 1) | NEW ENGLAND BioLabs | E7335S |
| Software and algorithms |  |  |
| Bowtie-1.0.0 | Langmead et al | https://genomebiology.biomedcentral.com/articles/10.1186/gb-2009-10-3-r25 |
| Samtools | Li and Durbin | https://academic.oup.com/bioinformatics/article/25/14/1754/225615 |
| MACS2 | Elizabeth et al | https://journals.plos.org/plosone/article?id=10.1371/journal.pone.0011471 |
| MiSeq System | Illumina | Cat# SY-410-1003 |
| S220 Focused-ultrasonicator | Covaris | Cat# 500217 |

### Synchronization and cell cycle analysis by Flow Cytometry

Yeast cells containing *nda3-KM311* mutation in PMG medium were arrested at 20 °C for 6 hr for cell cycle synchronization at M-phase. They were released into sub-sequential cell cycle at 30 °C. Cells in 5 mL culture were collected and suspended in 200 µL water. Cells were fixed with 600 µL ethanol, washed with 50 mM sodium citrate (pH7.5) (FUJIFILM Wako), and were treated with 0.1 mg/mL RNase A (Sigma-Aldrich) in 300 µL of 50 mM sodium citrate at 37 °C for 2 hr. Cells were stained with 4 ng/mL propidium iodide (Sigma-Aldrich) at room temperature for 1 hr. After sonication, cells were analyzed by BD LSRFortessaTM X-20.

### Co-immunoprecipitation

The procedure was performed as described previously (Shimmoto et al., 2009). For immunoprecipitation, approximately 1.0 ×10^8^ cells from 50 mL culture were harvested and washed once with PBS. The cells were then resuspended in 0.5 mL of IP buffer (20 mM HEPES-KOH [pH 7.6] (Nacalai tesque), 50 mM potassium acetate (Sigma-Aldrich), 5 mM magnesium acetate (FUJIFILM Wako), 0.1 M sorbitol (FUJIFILM Wako), 0.1% TritonX-100 (Sigma-Aldrich), 2 mM DTT (FUJIFILM Wako), 20 mM Na_3_VO_4_ (Sigma-Aldrich), 50 mM β-glycerophosphate (Sigma-Aldrich) and Protease Inhibitor Cocktail (Sigma-Aldrich)) and were disrupted with glass beads using a Multi-Beads Shocker (Yasui Kikai; Osaka, Japan). The lysates were cleared by centrifugation (15,000 rpm for 10 min at 4 °C). The supernatants of lysates were mixed with anti-c-Myc antibody (Nacalai tesque) attached to Protein G Dynabeads^TM^ (Thermo Fisher, 10004D). After incubation for 1 hr, beads were washed with IP buffer and proteins were extracted by boiling with 1× sample buffer (2% SDS (Nacalai tesque), 4 M Urea (Nacalai tesque), 60 mM Tris-HCl [pH6.8] (Nacalai tesque), 10% Glycerol (nacalai tesque), and 70 mM 2-Mercaptethanol (Sigma-Aldrich)].

### Immunoblot

Protein samples and prestained molecular weight markers (Bio-Rad) were loaded onto 5∼20% gradient precast PAGE gel (ATTO corp.), and transferred to PVDF membranes (Millipore). The membranes were blocked with 5% skim milk in TBST and target proteins were detected with ANTI-FLAG® M2 antibody (Sigma-Aldrich) and anti-***α***-tubulin (SantaCruz).

### Chromatin immunoprecipitation (ChIP)

1.0 ×10^9^ cells were cross-linked with 1% formaldehyde for 15 min at 30 °C, and prepared for ChIP as previously described (Y. Kanoh et al., 2015; Katou et al., 2003). Briefly, cross-linked cell lysates prepared by multi-beads shocker (Yasui Kikai Co.) and sonication (Branson) were incubated with Protein G Dynabeads^TM^ (Thermo Fisher, 10004D) attached to ANTI-FLAG^®^ M2 antibody (Sigma-Aldrich) for 4 hr at 4 °C. The beads were washed several times and the precipitated materials were eluted by incubation in elution buffer (50 mM Tris-HCl [pH7.6], 10 mM EDTA and 1 % SDS) for 20 min at 68 °C. The eluates were incubated at 68 °C overnight to reverse crosslinks and then treated with RNaseA (Sigma-Aldrich) and proteinase K (FUJIFILM Wako). DNA was precipitated with ethanol in the presence of glycogen (Nacalai tesque) and further purified by using QIAquick PCR purification kit (Qiagen).

### Living cell analysis

Cells were observed on BZ-X700 (KEYENCE) equipped with Nikon PlanApoλ 100× (NA=1.45) using IMMERSION OIL TYPE NF2 (Nikon). Mitotic spindles were visualized by expressing Pmt1-GFP-αTubulin. DNA damages were detected by observing fluorescent Rad52 foci (EGFP or YFP). Securin and spindle pole bodies were visualized by expressing Cut2-GFP and Sad1-GFP, respectively. The time-lapse images were observed on PMG medium/ 2% agarose (Nacalai tesque). Whole chromosome locations were visualized by expressing hht2 (Histone H3 h3.2)-GFP.

### Next generation sequencing (NGS) and ChIP-Seq

NGS libraries were prepared as described previously (Y. Kanoh et al., 2015). The input and the immunoprecipitated DNAs were fragmented to an average size of approximately150 bp by ultra-sonication (Covaris). The fragmented DNAs were end-repaired, ligated to sequencing adapters and amplified using NEBNext^®^ Ultra^TM^ II DNA Library Prep Kit for Illumina^®^ and NEBNext Multiplex Oligos for Illumina^®^ (New England Biolabs). The amplified DNA (around 275 bp in size) was sequenced on Illumina MiSeq to generate single reads of 100 bp. The generated ChIP or Input sequences were aligned to the *S. pombe* genomic reference sequence provided from PomBase by Bowtie 1.0.0 using default setting. Peaks were called with Model-based analysis of ChIP-Seq (MACS2.0.10) using following parameters; macs2 callpeak -t ChIP.sam -c Input.sam -f SAM -g 1.4e107 -n result_file –B -q 0.01. The pileup graphs were loaded on Affymetrix Integrated Genome Browser (IGB 8.0).

### In-gel kinase assay

In-gel kinase assays for replication checkpoint activation were conducted as described previously (Geahlen, Anostario, Low, & Harrison, 1986; Takeda et al., 2001; Waddell, Fialkow, Chan, Kishimoto, & Downey, 1995). SDS-polyacrylamide gel (10%) was cast in the presence of 0.5 mg/ml myelin basic protein (Sigma) within the gel. Extracts (100 µg of protein) prepared by the boiling method were run on the gel. After electrophoresis, the gel was washed successively in 50 mM Tris-HCl (pH 8.0), 50 mM Tris-HCl (pH 8.0)+5 mM 2-mercaptoethanol, and denatured in 6 M guanidium hydrochloride (Nacalai tesque) in 50 mM Tris-HCl (pH 8.0)+5 mM 2-mercaptoethanol, and renatured in 50 mM Tris, pH 8.0+5 mM 2-mercaptoethanol+0.04% Tween 20 over 12-18 h at 4°C. The gel was then equilibrated in the kinase buffer containing 40 mM HEPES-KOH (pH 7.6), 40 mM potassium glutamate, 5 mM magnesium acetate, 2 mM dithiothreitol, and 0.1 mM EGTA for 1 h at room temperature, and was incubated in the same kinase buffer containing 5 µ M ATP and 50 µCi of [γ -^32^P]ATP for 60 min at room temperature, followed by extensive washing in 5% trichloro-acetic acid (Nacalai tesque)+1% sodium pyrophosphate until no radioactivity is detected in the washing buffer. The gel was dried and autoradiographed.

### Cell fractionation and immunofluorescence analyses

5.0 × 10^7^ exponentially growing yeast cells were collected, and cell components were fractionated as previously reported (Y. Kanoh et al., 2015). Briefly, the cell walls were digested with 100 U/ml zymolyase (Nacalai Tesque) in 1.2 M sorbitol/ potassium phosphate (pH7.0) containing 1 mM PMSF at 30°C for 30 min. The spheroplast cells, washed with 1 M sorbitol, were permeabilized in a solution containing 0.1% Triton X-100 (Sigma-Aldrich), 1.2 M sorbitol/ potassium phosphate (pH 7.0) and 1 mM PMSF on ice. The cells were suspended in CSK buffer (50 mM HEPES-KOH [pH 7.5], 0.5% Triton X-100, 50 mM potassium acetate, 1 mM MgCl2, 1 mM EDTA, 1 mM EGTA, 1 mM DTT, 1 mM PMSF, 0.5 mM sodium orthovanadate, 50 mM NaF, 1× protease-inhibitor cocktail (Sigma-Aldrich), 1× protease-inhibitor cocktail (Roche), and 0.1 mM MG-132) for 30 min on ice. Genomic DNA was digested with 0.25 U/ml DNase I in CSK buffer containing 10 mM MgCl_2_ and 10 mM CaCl_2_ and incubated at 20°C for 30 min. The cells were fixed with 4% paraformaldehyde/ PBS after wash with CSK buffer. Nup98, a marker of the nuclear membrane, was detected with rat anti-Nup98 monoclonal antibody (1:500; Bioacademia) for 12 hr at 4°C after blocking in PBS containing 3% BSA and 0.1% Tween 20. The cells were washed with PBS containing 0.1% Tween 20 three times, and were incubated with Alexa Fluor 488– conjugated rabbit anti–rat IgG (1:1000; Invitrogen) in PBS containing 0.1% Tween 20 for 12 hr at 4°C. Antibodies were diluted in 1% BSA in PBS and 0.1% Tween 20.

Finally, the cells were stained with 1 µg/mL Hoechst 33342 for 1 hr at r.t. and washed with PBS containing 0.5% Tween 20 three times before visualization under microscope.

### Time laps analyses of cellular dynamics of Rif1

Cells expressing Rif1-mKO2 (red) and Taz1-EGFP (green) at the endogenous loci were analyzed under spinning disk microscope. Images were taken as previously reported (Ito et al., 2019) with slight modification. Briefly, microscope images were acquired using an iXon3 897 EMCCD camera (Andor) connected to Yokokawa CSU-W1 spinning-disc scan head (Yokokawa Electric Corporation) and an OlympusIX83 microscope (Olympus) with a UPlanSApo 100× NA 1.4 objective lens (Olympus) with laser illumination at 488 nm for GFP and 561 nm for mKO2. Images were captured and analyzed using MetaMorph Software (Molecular Devices). Optical section data (41 focal planes with 0.2 µm spacing every 2 min) were collected for 2 hr. Time-lapse images were deconvoluted using Huygens image analysis software (Scientific Volume Imaging).

---

**Supplementary Figure S1.**
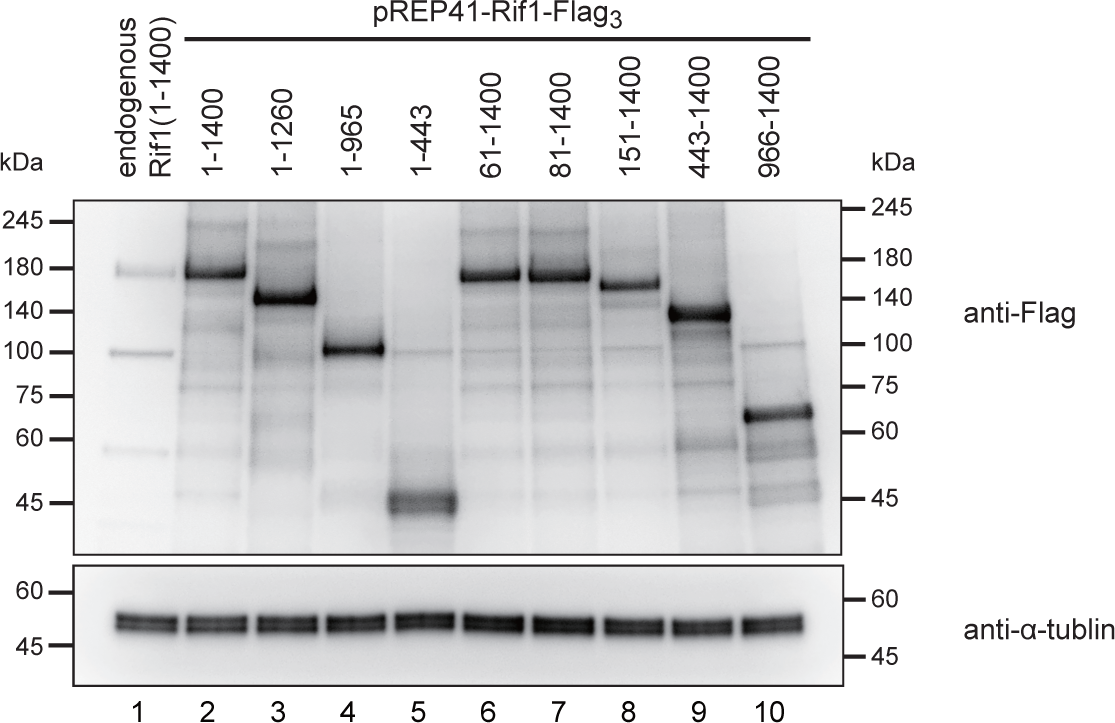
Expression levels of Rif1 and its derivatives. Western blot analyses of expression of the full-length Rif1 and its deletion/ truncation derivatives expressed on a plasmid after transfer to medium lacking thiamine for 24 hr. All the proteins carry His_6_-Flag_10_ tag at the C-terminus and proteins in the whole cell extracts were detected by anti-Flag antibody. ***α***-Tubulin protein level is shown as a loading control.

**Supplementary Figure S2.**
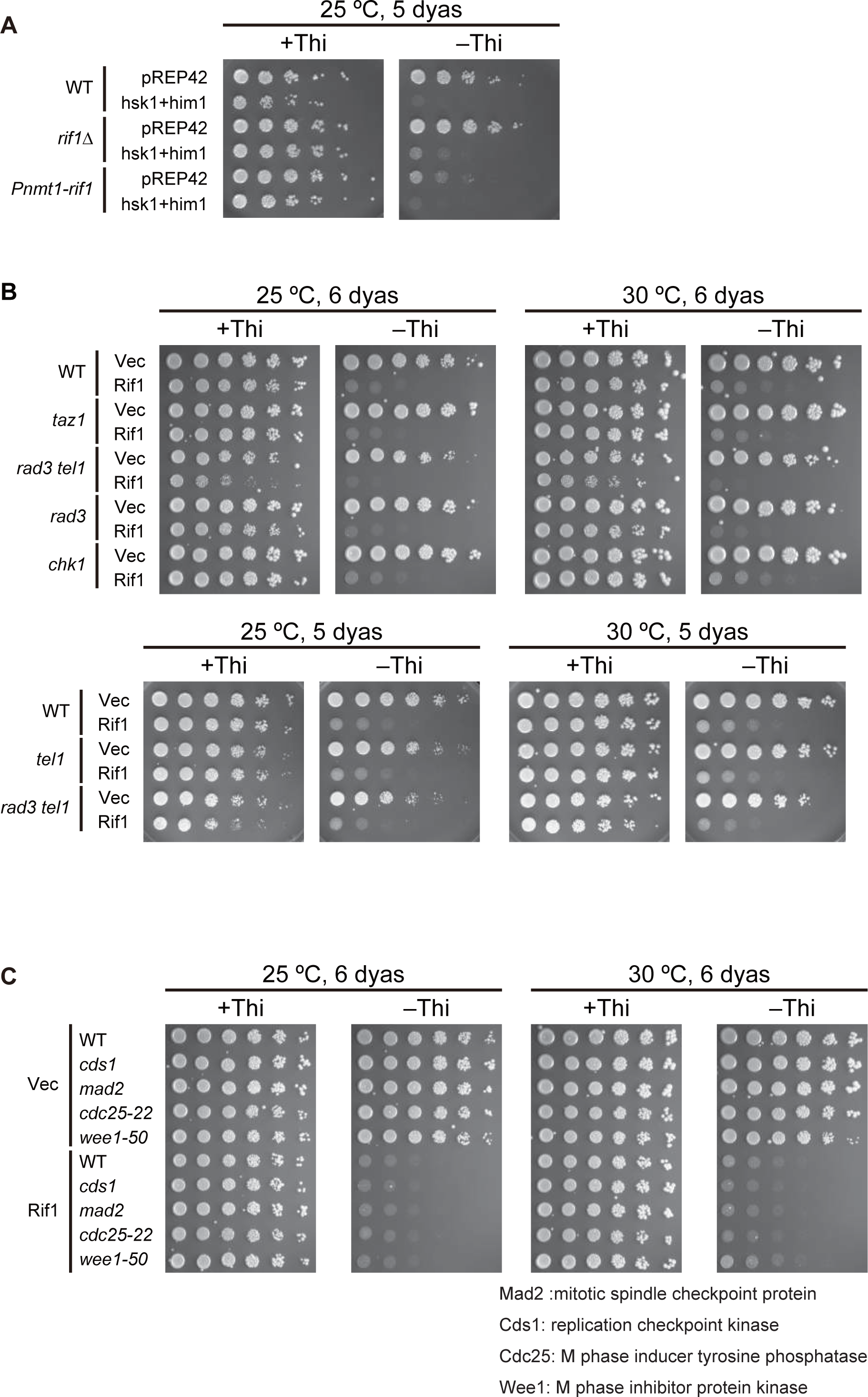
Effects of Rif1 overproduction on cell growth in various mutants. (**A**) Spot tests of the wild-type (WT), *rif1*Δ or Rif1-overproducing (Pnmt1-*rif1*) cells harboring vector (pREP42) or Hsk1+Dfp1/Him1 overproducing plasmid. (**B**) Spot tests of the wild-type (WT) and replication checkpoint mutant cells harboring vector (Vec) or Rif1-overproducing (Rif1) plasmid. (**C**) Spots test of the wild-type (WT) and various mutant cells harboring vector (Vec) or Rif1-overproducing (Rif1) plasmid. Proteins are overproduced on plates lacking thiamine (-Thi). Plates were incubated as indicated and photos were taken.

**Supplementary Figure S3.**
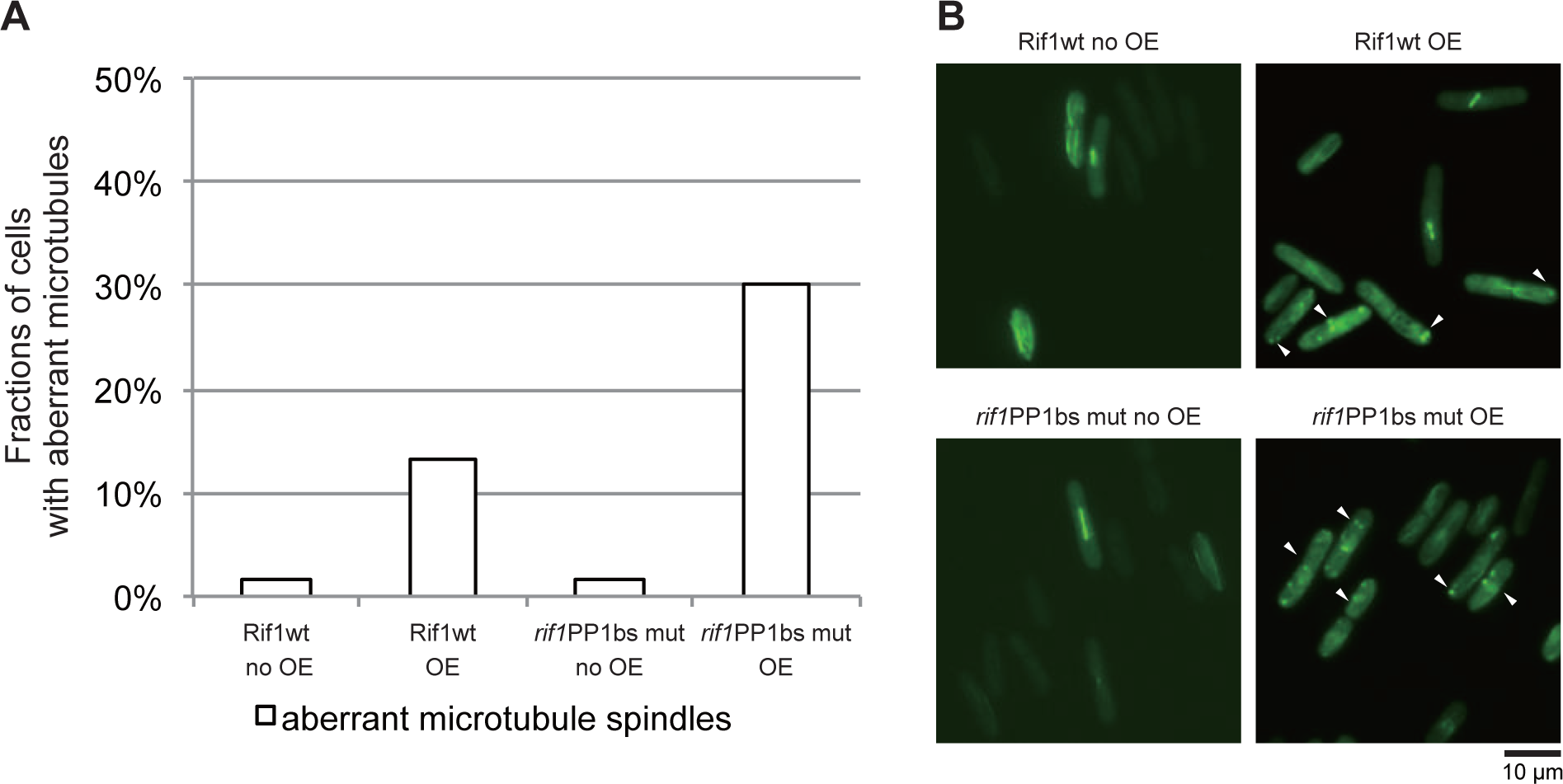
Cells with aberrant microtubules increase upon overproduction of Rif1. (**A**) Rif1 wt or PP1bs mutant was overexpressed in cells expressing GFP-α2tub and cells with aberrant microtubule spindles were counted. (**B**) The photos of cells with aberrant microtubule spindles are shown (indicated by arrowheads). In **A** and **B**, cells were grown in medium lacking thiamine for 18 hr.

**Supplementary Figure S4.**
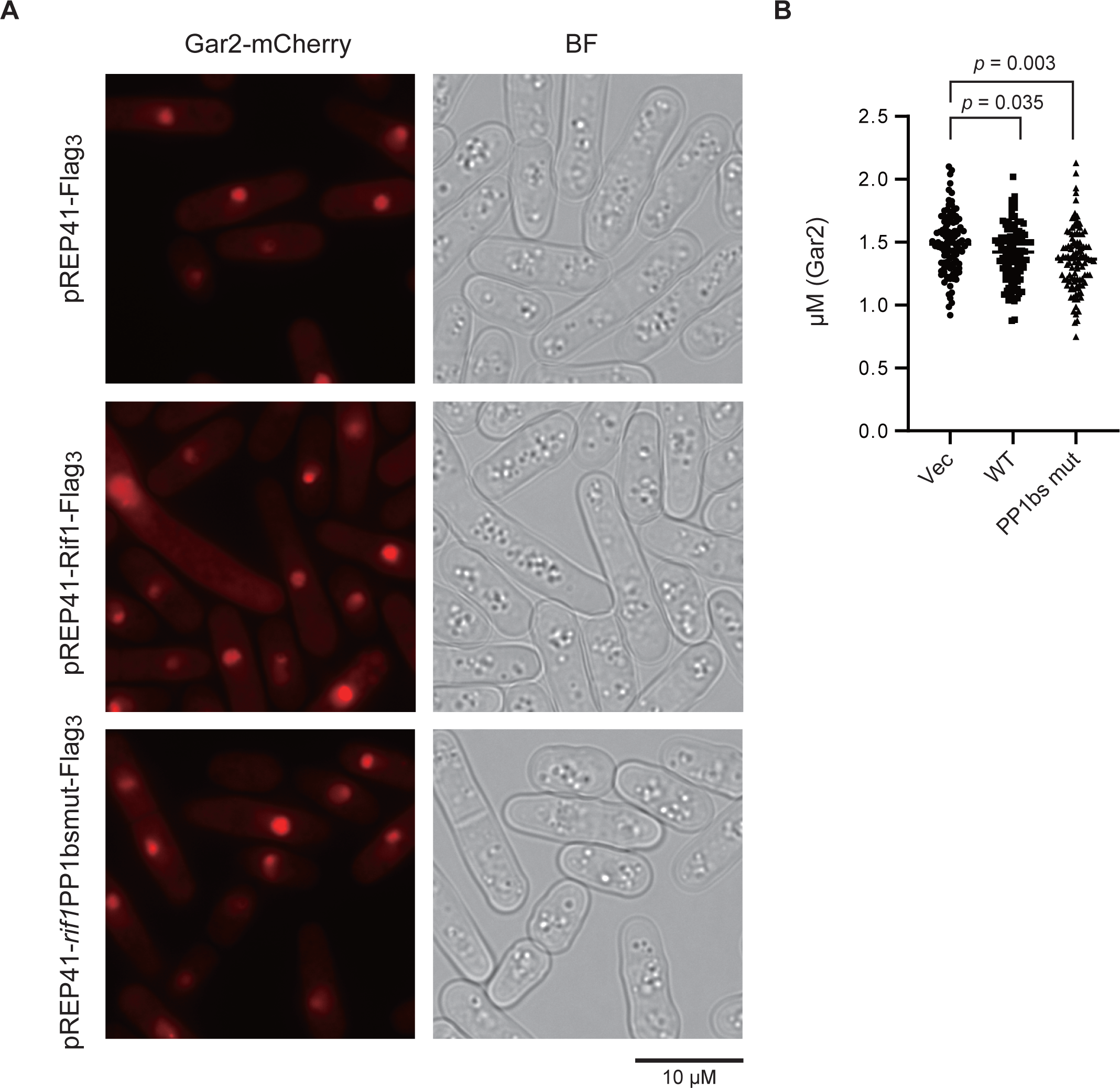
Sizes of nucleoli are not affected by overexpression of Rif1. **(A**) Rif1 wt (pREP41-Rif1-Flag_3_) or PP1bs mutant (pREP41-*rif1*PP1bsmut-Flag_3_) was overexpressed in cells expressing Gar2-mCherry (a marker for nucleoli) and the sizes of nucleoli were measured. pREP41-Flag_3_ represents the vector control. (**B**) The graph shows quantification of the data in (A). Y-axis shows the sizes of nucleoli, as measured by those of mCherry signals (diameter).

**Supplementary Figure S5.**
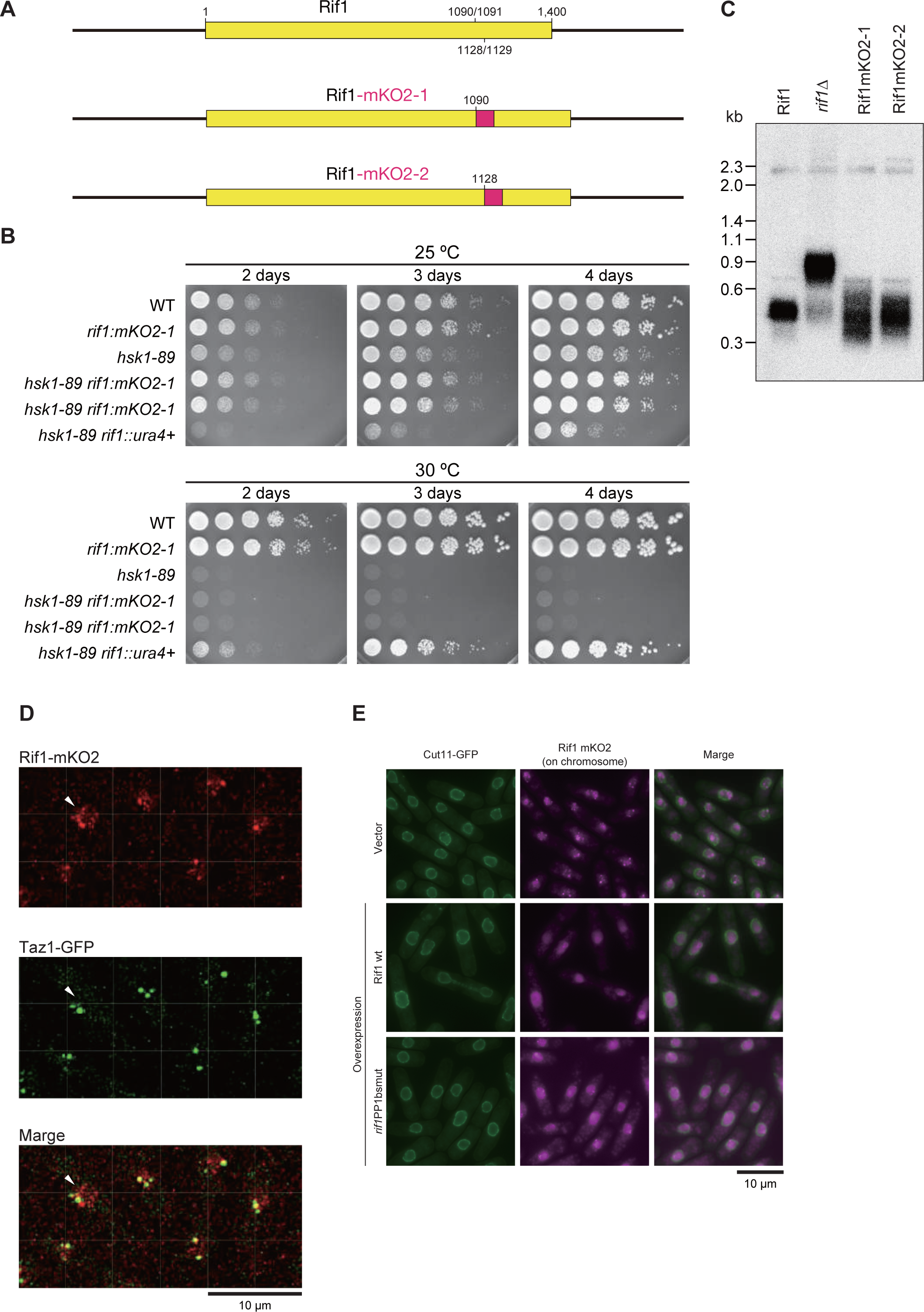
Evaluation of functions of Rif1-mKO2 fusion proteins. (**A**) Schematic drawing of Rif1-mKO2 fusions. mKO2 polypeptide was inserted at aa1090/1091 or at aa1128/1129, which is shown as a red box (not to the actual size). (**B**) Cells with indicated genotypes were serially diluted and spotted on EMM plates, and incubated at the indicated temperatures, as shown. Growth of *hsk1-89*(ts) at 30°C is not complemented by the Rif1-mKO2 fusions, suggesting they are functional. (**C**) Cellular DNA isolated from the strains shown were digested by *Eco*RI and probed by telomere-specific ^32^P-labebed DNA. (**D**) Enlarged images of Rif1-mKO2 (red) and Taz1-EGFP (green) taken from Movies 1∼3. The cells indicated by an arrow is focused in movie 5. In addition to several large Rif1-mKO2 foci that colocalize with Taz1, fine and dynamically moving Rif1-mKO2 foci, that are likely to represent Rif1 on chromosome arms, can be detected in nuclei. (**E**) Rif1-mKO2 signals (mazenta) in cells expressing Cut11-GFP (green). Cut11-GFP shows nuclear membrane. Upper panels, cells harboring a vector; middle panels, cells overexpressing wild-type Rif1 (pREP41-Rif1-Flag_3_); lower panels, cells overexpressing PP1bs mutant Rif1 (pREP41-*rif1*PP1bsmut-Flag_3_). Images were captured by KEYENCE BZ-X700 microscopy.

**Supplementary Figure S6.**
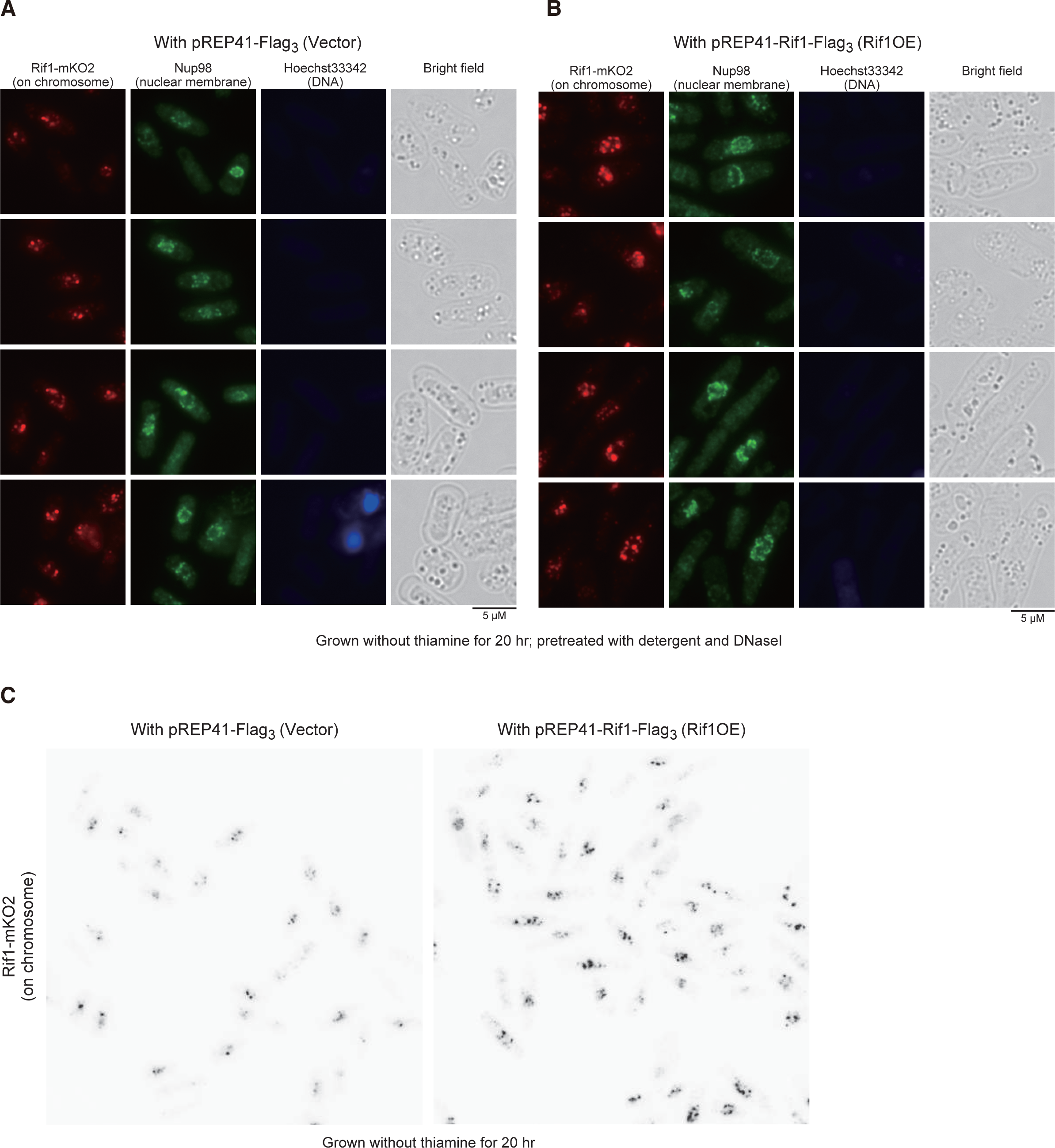
Effects of Rif1 overexpression on nuclear profiles of the endogenous Rif1 protein tagged with mKO2 (Rif1-mKO2). Rif1-mKO2 cells harboring vector (**A** and left panel of **C**) or Rif1-overexpressing plasmid (**B** and right panel of **C**) were pretreated with Triton X-100 and DNase I, and stained with anti-Nup98 antibody (green; nuclear membrane) and Hoechstm33342 (blue; nuclei). The Hoechst signal is very low due to prior treatment with DNase I. In **A** and **B**, mKO2 signals are in red, while they are in black in **C**.

## Supplementary movies

Cells expressing Rif1-mKO2 (red) and Taz1-EGFP (green) at the endogenous loci were analyzed under spinning disk microscope. Images were taken at every 2 min for 2 hr as described in “Materials and Methods”. The video presented is after deconvolution.

Movie 1 (Rif1-mKO2, red), movie 2 (Taz1-EGFP, green), and movie3 (red+green) are the maximum intensity projection of 3D image data in 2D space. Movie 4 is a 3D image reconstruction of an earliest time point in movies 1∼3. Movie 5 is an enlarged version of movie 3, focusing on the cell indicated in **Supplementary Fig. S5 D**.

